# Competition between co-localized gut symbionts underlies inter-individual variation in the honeybee gut microbiota

**DOI:** 10.64898/2026.07.09.737433

**Authors:** Gonçalo Santos-Matos, Jérôme Benedetti, Malick Ndiaye, Jana Počuča, Estelle Pignon, Shrihari Negi, Ryo Miyazaki, Yolanda Schaerli, Macarena Marín Arancibia, Philipp Engel

## Abstract

Microbes in the animal gut compete to colonize spatially restricted host niches. However, whether competition for space drives inter-individual variation among hosts, and which factors determine the outcome of such competition, remain poorly understood. Here, we investigated competitive interactions of the western honeybee gut symbiont *Frischella perrara* with other members of the bee gut microbiota. Shotgun metagenomics analysis of individual bees revealed that *F. perrara* is negatively correlated with a specific species of the genus *Gilliamella*. Co-colonization of microbiota-depleted bees with these two bacteria resulted in their competitive exclusion. The outcome of this competition depended on the relative number of bacteria each bee received and benefited *Gilliamella* when one of the two type VI secretion systems of *F. perrara* was mutated. Using fluorescently tagged strains, high-resolution microscopy, and gut region–specific quantification, we show that both bacteria localize to the same host niche in the ileum of mono-colonized bees, indicating competition in a spatially restricted host niche. Moreover, both microbes protected against infection, promoting bee health. This competition provides an explanatory mechanism underlying variation in the occurrence of *F. perrara* across honeybee colonies and highlights the importance of gut spatial structure and microbial competition in shaping microbiome composition and inter-individual variability.

## Introduction

Gut-associated bacterial communities play an important role in animal development, physiology, behaviour and overall fitness. Despite this importance, gut microbiota composition varies substantially between individuals of the same species. Such inter-individual variation has been linked to alterations in host health outcomes (1, 2). Elucidating the drivers of inter-individual variation in gut microbiota composition will advance our understanding of gut microbiota–host interactions and support the development of preventative and interventional approaches to modulate microbiota composition.

Besides host factors or diet consumption, bacterial competition is considered as a key factor in determining differences in community composition (3). It is well established that bacteria in the animal gut compete for shared nutrient resources, leading to competitive exclusion (4, 5). However, bacteria also compete for space, as colonization of a restricted niche within the gut can confer fitness advantages (6). Type VI secretion systems (T6SS) are microbial weapons that play an important role in spatial competition (7–10). These spear-like extracellular structures enable bacteria to inject toxins into neighboring cells in a contact-dependent manner (11, 12). Their widespread distribution among gut bacteria suggests that T6SS-mediated interference is a major driver of competition within this environment (13–16). Moreover, theory predicts that early numerical differences in the abundance of bacteria can determine the outcome of competition via interference, even if the inferior competitor has a higher intrinsic growth rate (17). Consequently, differences in the initial cell numbers of bacterial inoculation during community assembly can lead to substantial differences in community composition among individual hosts (17–19). Yet, little is known about competition for space in the animal gut, how inoculum size and T6SS modulate this competition, and if it can contribute to inter-individual variation among hosts. Addressing these questions is challenging given the complexity of animal gut ecosystems, which contain many spatial niches and exhibit high microbial diversity and variability.

The western honey bee *Apis mellifera* harbors a relatively simple and host-specific gut microbiota that is spatially structured along the hindgut (20, 21). While the posterior hindgut compartment, the rectum, is dominated by bacteria from the genera *Lactobacillus*, *Bombilactobacillus*, and *Bifidobacterium*, the anterior compartment, the ileum, primarily harbors species of the genera *Snodgrassella*, *Gilliamella*, and *Frischella* (22, 23). These proteobacteria form multispecies biofilms attached to the cuticular surface of the gut epithelium (24, 25) and are equipped with T6SSs (26), suggesting that they compete for host surface colonization via interference competition. *Frischella perrara*, the only species of the genus *Frischella* present in *A. mellifera* (27), predominantly colonizes the anterior section of the ileum, where it causes a localized host immune response and the formation of a characteristic brown-to-black deposit, referred to as the “scab” (25, 28). The prevalence of this bacterium and the occurrence of scab phenotypes vary greatly among honeybee colonies, with scabs detected in 15% to 80% of adult worker bees within a colony (25, 29). However, the ecological mechanisms underlying this striking variation in microbiota composition across conventional bees remain unknown. In a previous study, we examined how the colonization of *F. perrara* mutants for putative colonization factors was impaired by competition with a synthetic community (SynCom) composed of other honeybee gut symbionts (30). While the wild-type strain consistently colonized gnotobiotic bees, deletion of *hcp2*, a core component of one of the two type VI secretion systems (T6SS-2), caused a strong colonization defect specifically in the presence of the SynCom. Based on these findings, we hypothesize that *F. perrara* competes with other gut symbionts for colonization of its spatial niche in the anterior ileum, and that competition contributes to the inter-individual variation observed among honey bees in the environment.

Here, we show that *F. perrara* abundance is negatively correlated with that of a prevalent *Gilliamella* species in the bee gut both in nature and in the laboratory. Co-colonization experiments with strains of both species reveal that the outcome of this antagonistic interaction depends on their relative inoculum sizes and on the presence of a functional *F. perrara* T6SS-2. Moreover, both bacteria promoted honey bee health by lowering mortality caused by a bee entomopathogen. Finally, we show that both bacteria adhere to the host cuticular surface in the same region of the anterior ileum, indicating competition within a shared physical niche underlies inter-individual variation for *F. perrara* colonization.

## Results

### *F. perrara* and a specific species *of Gilliamella* are negatively correlated in the gut of honeybees

To identify candidate competitors of *F. perrara,* we examined correlations between its abundance and that of other honey bee gut symbionts in a) the SynCom experiment described above, in which microbiota-deprived (MD) bees were colonized with a synthetic community composed of the most frequent honeybee gut symbionts and different genotypes of *F. perrara* (30), and b) a gut metagenomic dataset consisting of wild-derived conventional adult worker bees of *A. mellifera* sampled from two hives in Switzerland (31) and two hives in Japan.

16S rRNA gene amplicon sequencing of the SynCom samples collected at day 10 post bacterial inoculation (n=80 bees) identified 13 ASVs corresponding to the six core genera included in the SynCom (**Supplementary Figure S1** and **Supplementary Figure S2**). We next determined the absolute abundance of each ASV by normalizing relative abundances to the total bacterial load of each sample, as determined by qPCR (see methods). Pairwise correlation analyses were then performed across all ASVs.

Among all taxa, a single ASV, *Gilliamella* ASV2, was negatively correlated with the ASV corresponding to *F. perrara* (ASV3, correlation coefficient: −0.29, p-value = 0.009) (**Figure 1A** and **1B**). In contrast, *Snodgrassella* ASV1, the other ileum colonizer, showed no correlation with *F. perrara* (correlation coefficient = −0.01, *p* = 0.94, **Figure 1A**).

**Figure 1.**
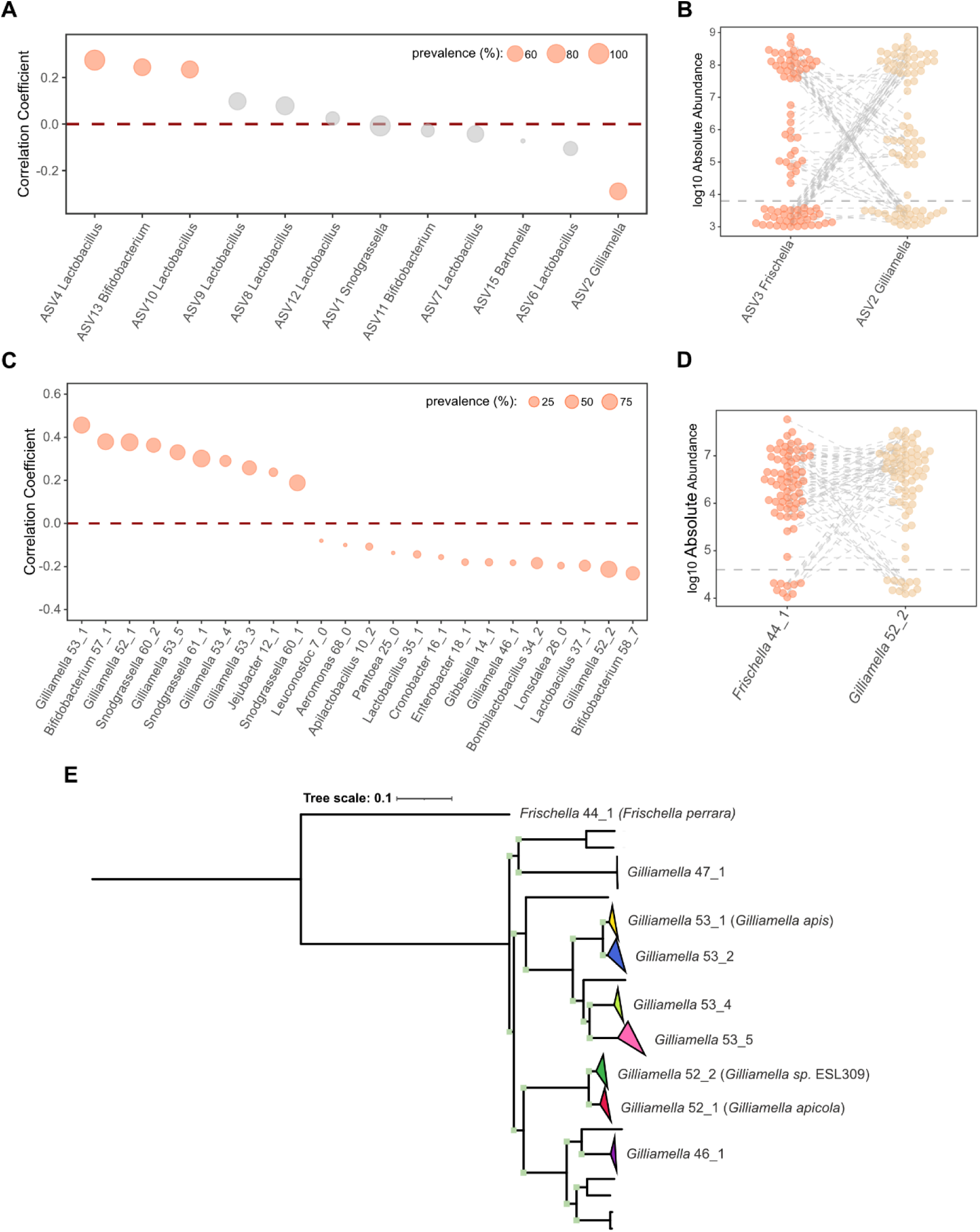
*F. perrara* is negatively correlated with a specific species of *Gilliamella*. **A** – Correlation coefficients (Y-axis) between the ASV corresponding to *F. perrara* and the remaining 12 ASVs (X-axis) identified by 16S rRNA gene amplicon sequencing on the samples of the SynCom experiment. Orange points represent significant correlations, while the size of the point highlights the prevalence of that ASV across the analyzed samples (n=80). **B** – Absolute abundance (Y-axis) of *Frischella* and *Gilliamella* ASVs (X-axis) in each SynCom sample. Orange and beige points represent *Frischella* loads and *Gilliamella* loads, respectively. The dotted grey lines connect values for the same bee sample. The horizontal grey dotted line represents the limit of detection, based on the number of reads and the total bacterial load in each sample. **C** - Correlation Coefficients (Y-axis) for all 24 species (X-axis) that had a significant correlation with *F. perrara* in the metagenomic analysis of gut samples from conventional *A. mellifera* (n=72 bees). **D** - Absolute abundance (Y-axis) of *F. perrara* 44_1 and *Gilliamella* sp. 52_2 (X-axis) in the metagenomic samples. Orange and beige points represent *F. perrara* 44_1 loads and *Gilliamella* 52_2 loads, respectively. The dotted grey lines connect bacterial loads for the same bee sample. The horizontal grey dotted line represents the limit of detection, based on the number of reads and the total bacterial load in each sample. **E** – Maximum-likelihood phylogeny of the *Gilliamella* genus based on the concatenated alignment of the protein sequences of 572 orthologous groups of 122 bacterial genomes (24 isolates and 98 metagenome-assembled genomes). Light green squares indicate bootstrap values above 90 (of 100 replicates). The LG+I+G4 substitution model with 1000 bootstrap replicates was used to calculate the phylogeny. Species names shown are based on the metagenomic analysis. *F. perrara* was used as an outgroup. For more details, see Supplementary Figure S6.

Independent quantification of the absolute abundance of these three taxa using genus-specific qPCR primers confirmed this pattern: in most samples and treatments where *F. perrara* successfully colonized, *Gilliamella* was undetectable, and *vice-versa,* while *Snodgrassella* loads were high across all tested conditions independent of the *F. perrara* loads (**Supplementary Figure S3**).

To test whether similar negative correlations occur in conventional honeybees (sampled from the natural environment), we used the same approach to analyse correlations in total bacterial abundances across the metagenomic samples (n= 73 bees). To this end metagenome-assembled genomes (MAGs) were established and clustered into 72 bacterial species (according to GTDB-TK), encompassing all major bee gut bacterial genera (**Supplementary Figure S4**). Among those, we identified a total of 24 significant correlations involving *F. perrara,* including ten with species of *Snodgrassella* and *Gilliamella* (**Figure 1C** and **Supplementary Figure S5)**. Eight of these ten correlations were positive. However, we identified two negative interactions of *F. perrara* with specific *Gilliamella* species, *Gilliamella* sp. 46_1 (correlation coefficient: −0.18, p-value = 0.048) and *Gilliamella sp.* 52_2 (correlation coefficient: −0.21, p-value = 0.042) (**Figure 1D** & **F)**. While *Gilliamella* sp. 46_1 was relatively rare and detected in only 4% of the samples, *Gilliamella sp.* 52_2 had a prevalence of 83%, much closer to that of *F. perrara* which colonized 86% of the analyzed honey bee samples (**Figure 1**, **Supplementary Figure S4)**.

To assess the phylogenetic position of *Gilliamella* sp. 52_2, we constructed a phylogeny of the genus *Gilliamella*, including the metagenome-assembled genomes from the metagenomic dataset as well as previously published genomes of isolate strains. *Gilliamella* sp. 52_2 belonged to a sister clade of *Gilliamella apicola* suggesting that it represents a distinct species within the genus *Gilliamella*. Notably strain ESL0309, one of the three *Gilliamella* strains included in the defined community used in the SynCom experiment, belonged to this same species and hence could be responsible for the observed negative correlation observed in this experiment (**Figure 1E** and **Supplementary Figure S6**). Altogether, these results suggest that *F. perrara* negatively interacts with a highly prevalent specific species of *Gilliamella*.

### Inoculum size determines the competitive outcome between *Frischella perrara* and *Gilliamella* sp. ESL0309 and is regulated by the type VI secretion system

To directly test whether *F. perrara* and *Gilliamella* sp. ESL0309 interact negatively, we inoculated MD bees with only these two bacteria. We used three inoculation ratios (1:10, 1:1, and 10:1) to test whether inoculum size determines the competitive outcome. Moreover, we competed *Gilliamella* sp. ESL0309 against either wild-type (wt) *F. perrara* or the T6SS mutant (Δhcp2) that showed a colonization effect in the SynCom experiment (30) to assess if the T6SS modulates the interaction outcome. To quantify the effect of competition on colonization, we compared the bacterial loads of both strains in the presence and the absence (monocolonization) of the competitor at day 10 post bacterial inoculation.

In competition with *F. perrara* wt, colonization by *Gilliamella* sp. ESL0309 was severely impaired when ESL0309 was either the less abundant strain (10:1) or inoculated at equal abundance (1:1), as indicated by significant 2-log fold changes in bacterial loads relative to monocolonization (**Figure 2A** and **Supplementary Figure S7**; 1:10, p-value < 0.001; 1:1, p-value = 0.001). When *Gilliamella* sp. ESL0309 was the more abundant strain in the inoculum (1:10), its colonization level was comparable to that observed in monocolonization (**Figure 2A** and **Supplementary Figure S7**; 1: 10, p-value = 0.338). In contrast, colonization of *F. perrara* was affected by the presence of *Gilliamella* sp. ESL0309 only when *F. perrara* was the less abundant strain at inoculation (**Figure 2A** and **Supplementary Figure S7**; 10:1, p-value = 0.525; 1:1, p-value = 0.284; 1:10, p-value = 0.025). Together, these results indicate that the two bacteria negatively impact each other’s colonization and that inoculum size determines the competitive outcome. Moreover, *F. perrara* was the more competitive strain when both species were introduced at equal abundance.

**Figure 2.**
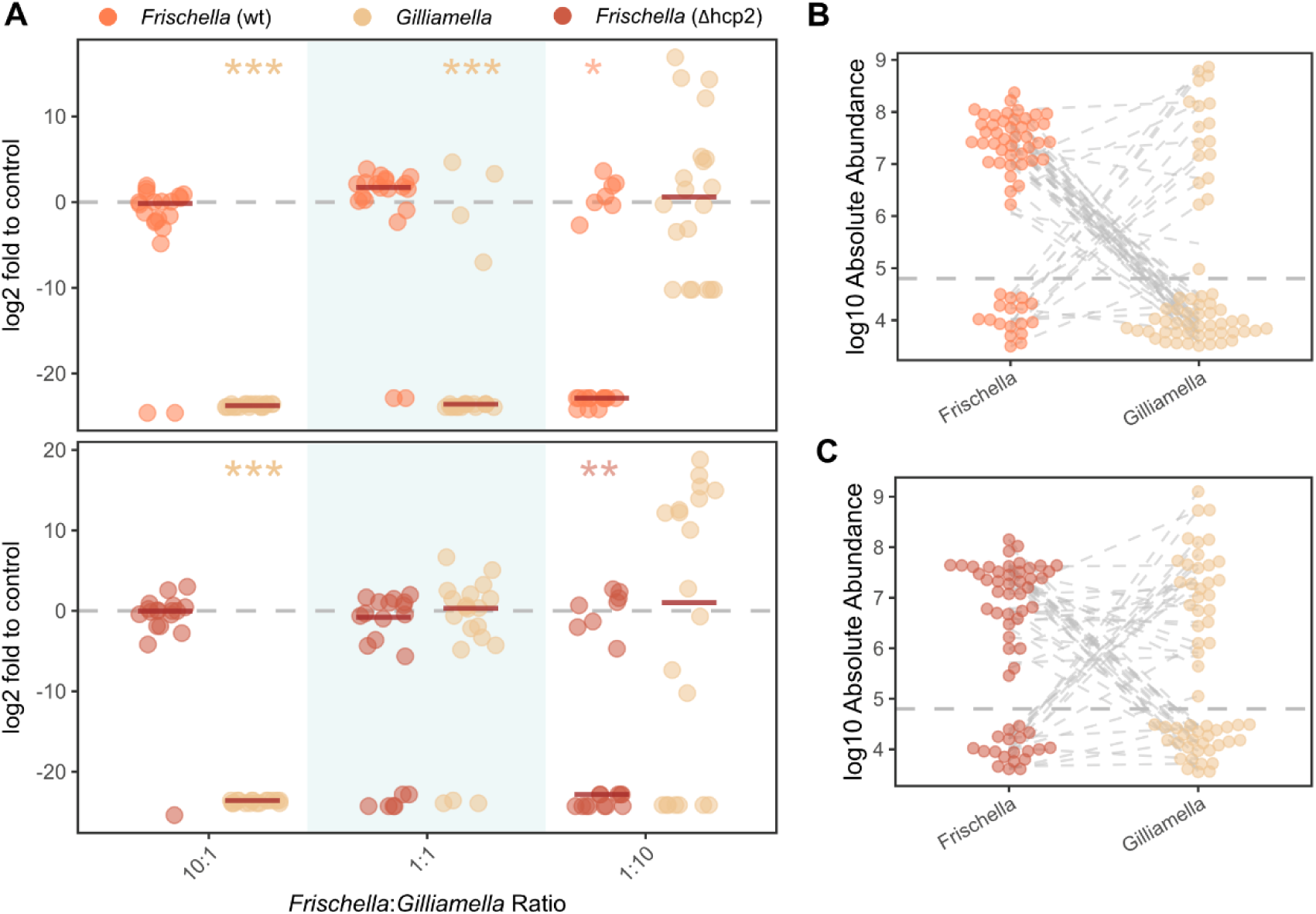
Inoculum size determines the competitive outcome between *F. perrara* and *Gilliamella* sp. ESL309 and is regulated by the type VI secretion system of *F. perrara*. A – Competition between *F. perrara* wt and *Gilliamella* sp. ESL309. and between *F. perrara* Δhcp2 and *Gilliamella* sp. ESL309. Bacterial loads represented on the Y-axis were obtained by calculating the log2 difference between bacterial loads in competition and the corresponding control in monocolonization, obtained using qPCR with genus specific primers. The three different inoculums tested are shown on the X-axis. Statistical analyses were based on pairwise Wilcoxon tests based on the colonization values prior to log2 calculation between competition and monocolonization. * p-value < 0.05, ** p-value < 0.01, *** p-value < 0.001. **B** – *F. perrara* wt and *Gilliamella* sp. 309 loads at day 10 post-inoculation for the samples belonging to the 10:1, 1:1 and 1:10 ratios altogether. **C** – *F. perrara* Δhcp2 and *Gilliamella* sp. 309 loads at day 10 post-inoculation for the samples belonging to the 10:1, 1:1 and 1:10 ratios altogether. For B and C, bacterial loads were quantified using genus specific primers and normalized to the host actin values. The diagonal dotted lines unite values belonging to the same honey bee sample. The horizontal line depicts the limit of detection based on the genus specific qPCR primers. Further information on the mono colonization controls (without competition) and bacterial loads discriminated by ratio and *F. perrara* genotype can be found in the **Supplementary Figure 7**.

This pattern changed when the T6SS mutant of *F. perrara* (Δhcp2) was competed against *Gilliamella* sp. ESL0309. While the outcomes of competition at the 1:10 and 10:1 inoculation ratios were comparable to those observed with *F. perrara* wt (**Figure 2A** and **Supplementary Figure S7**), colonization by *Gilliamella* sp. ESL0309 was no longer impaired when *F. perrara* Δhcp2 was introduced at equal abundance (1:1, p-value = 0.843). These findings demonstrate that the T6SS-2 of *F. perrara* modulates the outcome of competition with *Gilliamella* sp. ESL0309. Notably, in many bees only one of the two bacteria managed to establish detectable colonization suggesting that the two bacteria often completely exclude each other. Only in the case when *F. perrara* Δhcp2 was inoculated at the same abundance as *Gilliamella* sp. ESL0309, we observed a large number of bees in which both bacteria could coexist (**Figure 2B & C, Supplementary Figure S8**). Altogether, these results show that inoculum size determines the competitive outcome between *F. perrara* and *Gilliamella* sp. ESL0309 and that this outcome is regulated by the T6SS-2.

### *F. perrara* and *Gilliamella* sp. ESL309 occupy a similar physical niche within the gut

Given the canonical mode of action of T6SSs is contact-dependent and that the *F. perrara* T6SS-2 mediates the exclusion of *Gilliamella* sp. ESL0309, we hypothesized these two microbes compete within the same physical niche of the honeybee gut. To address this, we colonized MD bees with either *F. perrara* wt or *Gilliamella* sp. ESL309. Seven days post colonization, the ileum region of the gut was dissected, separated into anterior and posterior portions and CFU counting was used to quantify bacterial loads. Full-length ileums were used as controls. We observed that, for both bacteria, loads in the anterior portion of the ileum did not differ from those found in the full ileum (*F perrara*: p-value = 0.109; *Gilliamella* sp.: p-value = 0.167) but were statistically higher than the bacterial loads in the posterior portion of the ileum (*F. perrara*: p-value=0.005; *Gilliamella* sp.: p-value = 0.001) (**Figure 3A**). This result suggests both bacteria preferentially colonize the anterior portion of the ileum, which is in line with what has been published previously for *F. perrara* (25, 30) and suggests that *Gilliamella* sp. ESL309 colonizes the same region.

**Figure 3.**
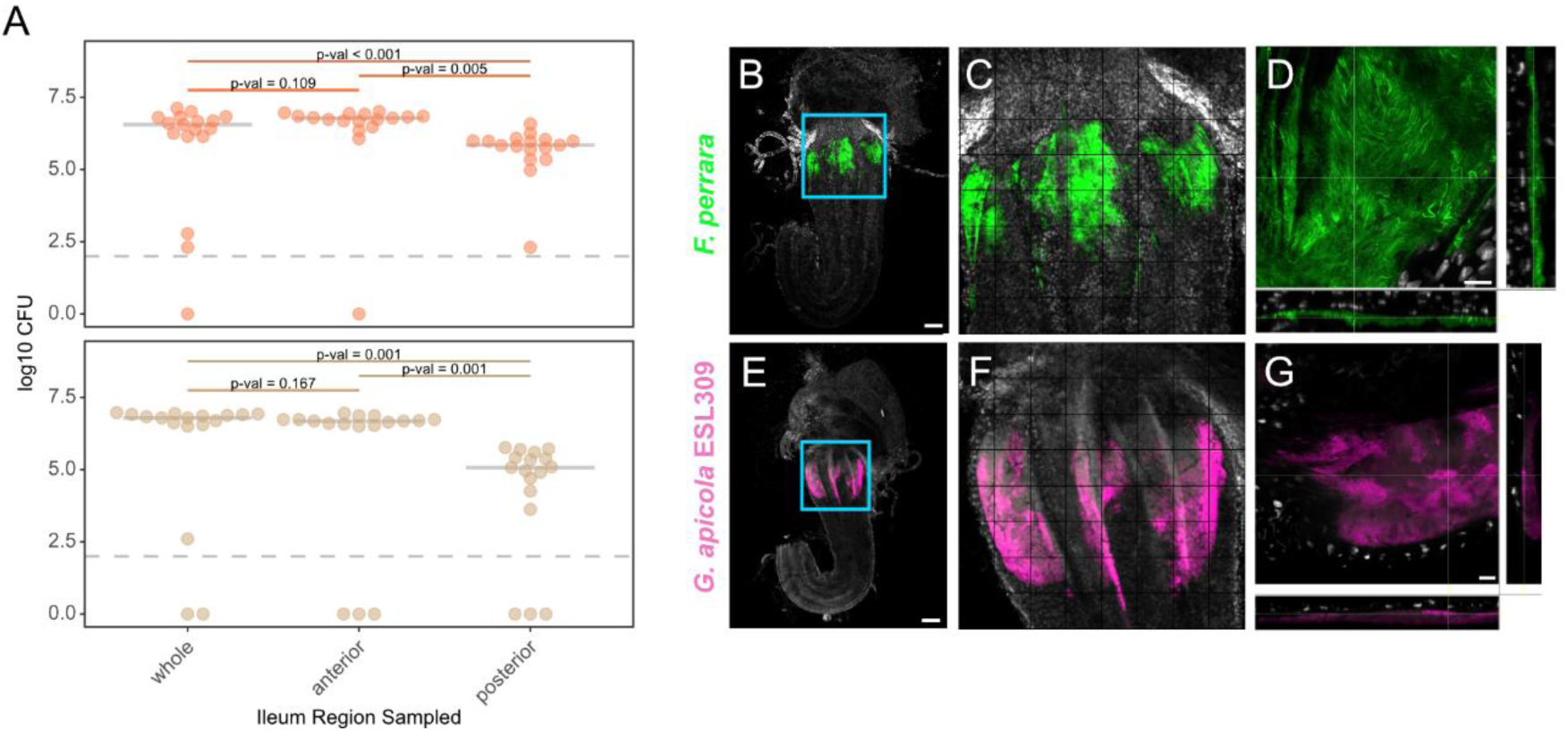
*F. perrara* and *Gilliamella sp.* ESL309 occupy a similar physical space within the anterior ileum. **A** – Microbiota-deprived bees were colonized with either *F. perrara* wt or *Gilliamella* sp. ESL309 and at day 7 post colonization the ileum was dissected, kept as a whole, or separated in anterior and posterior part, and bacterial abundances were obtained by CFU quantification. **B** – Microbiota-deprived bees were colonized with *F. perrara* engineered with the pAC08 plasmid expressing GFP and at day 7 post colonization, the anterior portion of the ileum was dissected and bacterial colonization was visualized using a Leica Stellaris Confocal microscope. **C –** Zoom in for the pylorus region of image B. **D** – Detailed visualization of *F. perrara* colonization of a crypt; panel on the bottom shows the 3D distribution of *F. perrara* on the X-axis while the panel on the right show the 3D distribution on the Y-axis. **E –** Microbiota-deprived bees were colonized with *Gilliamella sp.* ESL309 engineered with the pAC09 plasmid expressing Crimson and at day 7 post colonization, the anterior portion of the ileum was dissected and bacterial colonization was visualized using a Leica Stellaris Confocal microscope. **F** – Zoom in for the pylorus region of image D. G - Detailed visualization of *Gilliamella sp.* ESL309 colonization of a crypt; panel on the bottom shows the 3D distribution of *F. perrara* on the X-axis while the panel on the right show the 3D distribution on the Y-axis. For images B to G, DAPI staining was used and shown with the white color. The scale bar for B and E corresponds to 200 µm and that for D and G correspond to 20 µm. Figure C and F are enlarged versions of B and E and include a grid on top to better visualize bacterial colonization and allow comparison between the two guts. Per gut, the grid size was normalized to the maximum length measured in the pylorus.

To visualize the detailed localization of the two bacteria, we generated fluorescently labelled strains of *F. perrara* wt (GFP) and *Gilliamella* sp. ESL309 (Crimson) (**Supplementary Figure S9**) and colonized MD bees with either of the two strains. As previously observed, *F. perrara* preferentially colonized the pylorus region (**Figure 3B and C**, **Supplementary Figure S10**). *Gilliamella* sp. ESL309 was also found within the pylorus and its distribution greatly overlaps with that of *F. perrara* (**Figure 3E & F; Supplementary Figure S10 and S11**). Visualization of colonized guts at increased magnification identified both microbes colonizing the crypts of the pylorus by forming compact biofilms (**Figure 3D & G**). These results reveal an overlap in the physical niches of *F. perrara* and *Gilliamella* sp. ESL309 within the anterior ileum.

### Both *F. perrara* and *Gilliamella* sp. ESL309 lower *Serratia*-induced mortality

Given that *F. perrara* induces a localized host immune response leading to the scab phenotype, whereas *Gilliamella* ESL0309 does not, we asked whether colonization by either bacterium differentially affects bee health. To test this, we measured survival of gnotobiotic honey bees inoculated with each bacterium alone and the two bacteria together (competition) relative to MD (i.e. microbiota-depleted) bees. Overall, there were no differences across treatments (p-value = 0.078). Nevertheless, contrast analysis revealed that honey bees inoculated with *Gilliamella* sp. ESL0309 survived significantly more than MD bees (p-value = 0.008 for the comparison using emmeans on the coxme model), while no such difference was found for the *F. perrara* (p-value = 0.785) and the co-colonization treatments relative to MD bees (p-value = 0.734) (**Figure 4 A & C)**.

**Figure 4.**
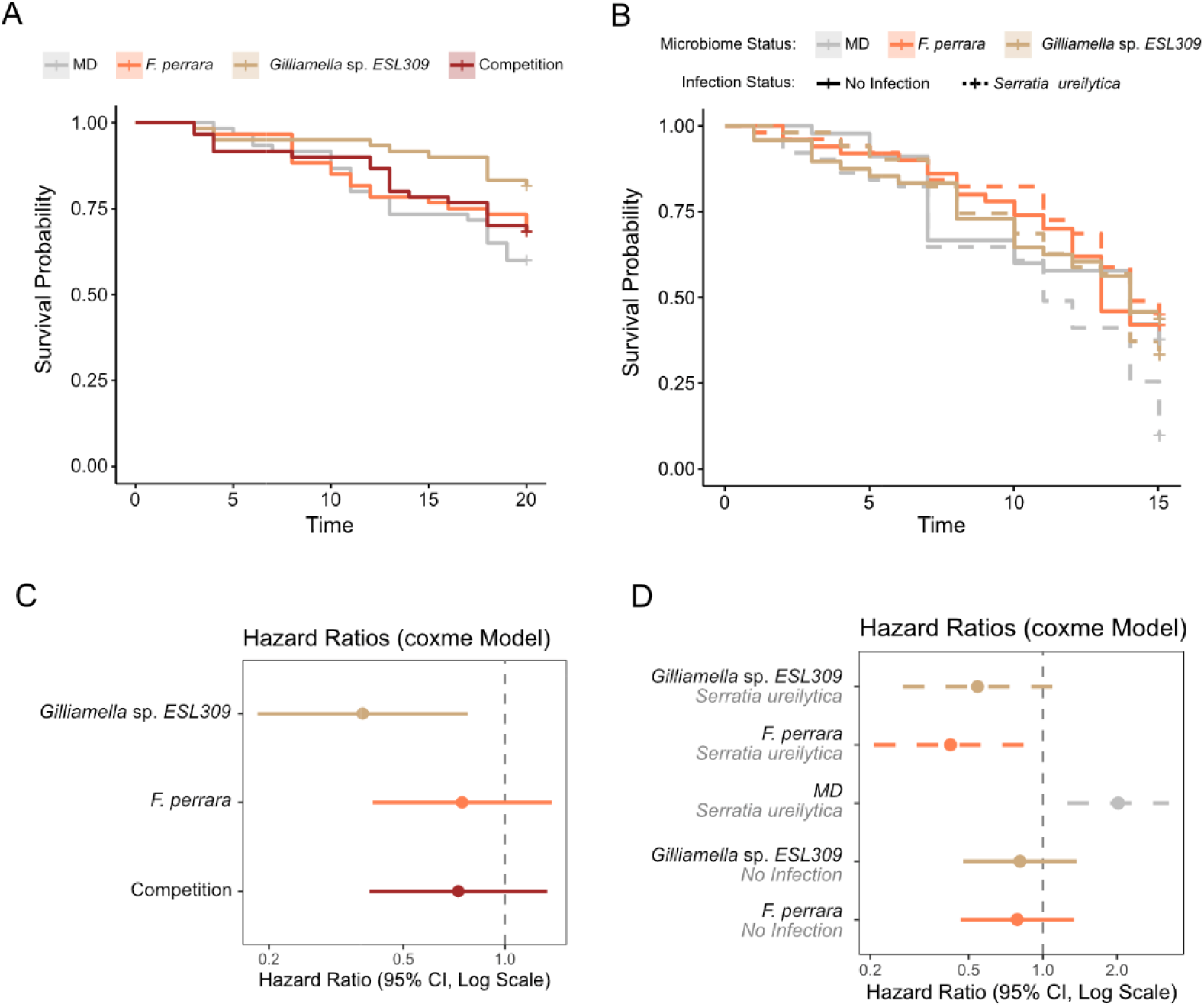
*F. perrara* and *Gilliamella* sp. ESL309 improve survival following infection with the entomopathogenic gut bacterium *Serratia ureilytica*. A – Survival curves of honey bees inoculated with *F. perrara*, *Gilliamella* sp ESL309, both bacteria simultaneously (competition) and no bacteria (MD). **B** – Honey bees were initially colonized with either *F. perrara*, *Gilliamella* sp ESL309 or not inoculated. 5 days after the first inoculation, bees were inoculated with *S. ureilytica* or PBS and survival post infection was measured. For both A and B, survival was monitored daily. **C** – Hazard ratios for the growth curves shown in A. **D** – Hazard ratios for the survival curves shown in B. Results obtained from 2 independent experiments, each containing 3 technical replicates per condition.

Then, we asked if *F. perrara* and *Gilliamella* sp. ESL0309 could improve survival following infection with the entomopathogenic gut bacterium *Serratia ureilytica* (32, 33). MD bees were inoculated with either *F. perrara*, *Gilliamella* sp. ESL0309, or no bacteria and 5 days later challenged by inoculating *S. ureilytica* or with a saline solution. Both *F. perrara* and *Gilliamella* sp. ESL0309 increased survivorship of bees infected with *S. ureilytica* relative to MD bees (p-value = 0.002 for *F. perrara*; p = 0.0061 for *Gilliamella* sp. ESL0309, results obtained using emmeans on the coxme model). There was no difference in the level of protection the two bacteria provided (p-value = 0.9856 for the comparison using emmeans on the coxme model) (**Figure 4 B & D)**. These results demonstrate that both microbes can promote honey bee health by conferring protection to a pathogen infection.

## Discussion

Bacteria compete for space and nutrients, yet how such competition contributes to inter-individual variation in gut communities remains poorly explored. Our data shows that *F. perrara* and *Gilliamella* sp. ESL309 are negatively correlated in conventional bees, inhibit each other’s colonization when co-inoculated into microbiota deprived bees, protect bees against infection and share a similar physical niche in the anterior ileum. The outcome of this competition depends on inoculation size and on the presence of a functional type VI secretion system (T6SS) in *F. perrara*, suggesting that their reciprocal exclusion is driven by interference competition. This mechanism provides a plausible explanation for why *F. perrara* is found only in a subset of bees within each colony and emphasizes the importance of competition between symbionts that occupy the same spatial niche in gut microbial communities.

Previous studies have shown that closely related bacterial strains and species coexist in the honey bee gut through the partitioning of metabolic and spatial niches. Such niche differentiation is thought to have been a key driver of diversification in several dominant bee gut symbiont lineages, including *Gilliamella* (24, 34, 35). In contrast, we show that two distantly related species have converged on a largely overlapping ecological niche. These microbes engage in strong competitive interactions that can lead to reciprocal exclusion, providing a potential mechanism underlying variation in gut microbiota composition in *Apis mellifera* populations, and also evolutionary changes in composition across other bee species.

Our metagenomic analyses revealed that the negative association between *F. perrara* and *Gilliamella* is species-specific: only two of the identified *Gilliamella* species were negatively correlated with *F. perrara*. Somewhat counterintuitively, most other *Gilliamella* species, as well as several *Snodgrassella* species, showed positive correlations with *F. perrara*. Given such samples contained the full hindgut (ileum and rectum), these positive associations may be driven by indirect factors, such as variation in the contribution of ileum versus rectum DNA to the sequencing data, rather than by direct facilitative interactions or enhanced colonization capacity. Moreover, since *Gilliamella* species localize to distinct regions of the ileum and exhibit different metabolic capabilities (24), some species may have reduced niche overlap with *F. perrara*. A similar argument applies to *S. alvi,* which colonizes the host epithelium throughout the ileum rather than being restricted to the anterior part (24, 25, 36) and relies primarily on organic acids rather than sugars as an energy source (37). Consistent with their distinct spatial distributions and resource utilization patterns, *S. alvi* colonization was unaffected by the presence of *F. perrara* in our experiments (**Supplementary Figures S1 & S3**), suggesting limited niche overlap between the two species. In contrast, our experiments confirmed a negative interaction between *F. perrara* and one of the two *Gilliamella* species that was negatively associated with *F. perrara* in our ecological survey of conventional bees. A similar antagonistic interaction was recently reported for a different strain of the same *Gilliamella* species (38), indicating that this relationship can be recapitulated experimentally and may constitute a consistent feature of their association.

A striking observation was that in gnotobiotic bees, *F. perrara* and *Gilliamella* sp. ESL309 often completely excluded each other from the bee gut, despite the fact that there was no competition with other bacteria. These results suggest that competition can contribute to inter-individual variation in gut microbiota composition. According to ecological theory, such variation arises when one competitor gains a colonization advantage early during community assembly, independent of intrinsic growth rates—for example, by arriving first in the gut or being present at higher initial abundance (18). Our results align with this as *F. perrara* excluded *Gilliamella* sp. ESL309 when more abundant at inoculation (10:1 ratio) but was outcompeted when less abundant (1:10 ratio) (**Figure 2**). Moreover, theory predicts that the benefits of interference scale with population size (17), which is supported by the observation that the T6SS-2 of *F. perrara* becomes more relevant as the *F. perrara* cell numbers increase relative to those of *Gilliamella* (**Figure 2 and Supplementary Figure S7**: 1:10 ratio compared to 1:1). In the hive, newly emerged adult bees acquire bacteria from contacts with multiple conspecifics and given *F. perrara* and *Gilliamella sp.* abundance varies between individuals (**Figure 1 Supplement 4, *Frischella* 44_1** and ***Gilliamella* 52_2)**, bees are likely exposed to different inoculation sizes, shaping symbiont colonization.

Honey bees colonized with *F. perrara, Gilliamella* sp. ESL309 or both bacteria simultaneously, survived equally well (**Figure 4A** & **C**). Moreover both bacteria conferred similar levels of protection against pathogen infection with *Serratia* (**Figure 4B** & **D)** (38). These results suggest that variation in the abundance or presence of these two bacterial taxa in the gut has no consequences for host survival following infection with this pathogen. A recent study suggested that *Frischella* provides protection against *Serratia* through colibactin-induced phage induction (38). *Gilliamella* does not encode this gene cluster, indicating that any protective effects it may confer are likely mediated by different mechanisms. Moreover, defense against pathogens is only one of the axes in which these bacteria may differentially influence their host. For example, bacteria of the genus *Gilliamella* have been shown to modulate host learning and memory (39). Whether *F. perrara* exerts similar effects on the host remains unknown. Thus, although no differences in pathogen protection were observed, variation in the abundance of these symbionts may nonetheless affect other host traits.

A previous study focusing on *Snodgrassella* showed that the T6SS contributes to intraspecific interactions among *Snodgrassella* strains but does not appear to mediate interspecific interactions with other members of the bee gut microbiota (40). In contrast, our findings indicate that the *F. perrara* T6SS is involved in antagonistic interactions across species boundaries targeting a member of a different genus. However, whether *F. perrara* directly delivers T6SS effectors into *Gilliamella* cells remains to be determined. Notably, the T6SS-deficient mutant of *F. perrara* fails to induce the characteristic scab phenotype on the bee gut epithelium (30). Similarly, the study on *Snodgrassella* showed that the T6SS can modulate the expression of host immune effectors (40). Therefore, although our results are consistent with a T6SS-dependent antagonistic interaction, it remains unclear whether this effect is mediated by direct intoxication of *Gilliamella* cells or indirectly through T6SS-induced changes in the host environment. Independently of the underlying mechanism, our findings suggest that the reciprocal exclusion of the two species is driven, at least partially, by competition for spatial niches. Microscopy analyses showed that both species are restricted to the epithelial invaginations in the anterior ileum, even in the absence of bacterial competition elsewhere in the ileum. Moreover, the competitive interaction between *F. perrara* and *Gilliamella* sp. ESL0309 was mediated via T6SS, a contact-dependent antagonistic mechanism, supporting the idea that competition for physical space contributes to their mutual exclusion. Previous studies have shown that *Gilliamella* and *Frischella* differ substantially in their metabolic repertoires, although they are both saccharolytic fermenters (41). Also, even closely related *Gilliamella* strains share only part of their metabolic potential (24, 35). Accordingly, competition for nutritional resources is unlikely to fully explain the observed antagonism. This interpretation is further supported by the fact that strong antagonistic interactions persisted despite unrestricted access to pollen and sugar water and the absence of other gut microbiota members in our experiments.

However, our results also indicated that the spatial niche is not completely overlapping as *F. perrara* sometimes colonized a smaller range of the anterior ileum than *Gilliamella* sp. ESL309 **(Supplementary Figure 10 & 11)**. Moreover, while in gnotobiotic bees the two bacteria often excluded each other, frequent co-occurrence was observed in conventional bees and in the laboratory condition in which *F. perrara* lacked its T6SS. Understanding the spatial-temporal dynamics of the colonization of this niche will provide relevant mechanistic insights into the processes underlying co-occurrence or complete exclusion. We hypothesize that if initial symbiont populations are sufficiently small, they will colonize different micro-niches (e.g. separate crypts) of the anterior ileum resulting in isolated growth and co-occurrence. Such patterns have been observed for the colonization of the stomach glands by *Helicobacter pylori* (42). Characterizing the spatial niches of more strains and species of *Gilliamella*, and testing how colonization is impaired by *F. perrara* will further contribute to our understanding of how spatial niche overlap underlies competition. Finally, additional mechanisms may contribute to the antagonism between these two species, including competition for specific metabolites or the activation of host immune pathways that impair the competitor’s ability to establish successful colonization, particularly when it is the inferior competitor.

Overall, our findings support the idea that inoculation size and spatial constraints interact to generate alternative community states in the gut. Together, our results underscore the importance of considering spatial structure and microbe-microbe interactions when interpreting inter-individual variability in host-associated microbial communities.

## Methods

### Bacterial strains and manipulation

Before each experiment, bacterial cultures kept at −80°C in 25% glycerol stocks were amplified in petri dishes containing solid media, and grown for 3-5 days in anaerobic conditions (8% H2, 20% N2, 78% CO2 in a Vinyl Anaerobic Chamber, Coy Lab) before further manipulation. While *F. perrara* strain PEB0191 (43) was grown in plates containing modified Tryptone Yeast extract glucose media (0.2% Bacto tryptone, 0.1% Bacto yeast extract, 2.2 mM D-glucose, 3.2 mM L-cysteine, 2.9 mM cellobiose, 5.8 mM vitamin K, 1.4 µM FeSO4, 72.1 µM CaCl2, 0.08 mM MgSO4, 4.8 mM NaHCO3, 1.36 mM NaCl, 1.8 µM hematine in 0.2 mM histidine, 1.25% Agar adjusted to pH 7.2 with potassium phosphate buffer), *Gilliamella* sp. ESL309 was amplified in Brain-Heart Infusion (BHI) petri dishes.

### Generation of gnotobiotic honey bees

Microbiota-deprived (MD) honeybees were obtained from *Apis mellifera carnica* colonies maintained at the University of Lausanne, as previously described (44). For experimental colonization, bees were starved for 1–3 hr by removal of the sugar water solution. Then, bees were cooled down to 4°C in a refrigerator or on ice to transfer them (head side first) into 1.5 ml microfuge tubes with a hole at the bottom. Tubes with bees were kept at room temperature (RT). For inoculation, each bee was fed 5 µl of *F. perrara* resuspended in a solution of sugar water and PBS (1:1 v/v) through the hole at the bottom of the microcentrifuge tube. The bacterial inoculums were adjusted to an OD_600_ of 0.01 or 0.1 depending on the experiment. Colonized bees were kept at 30°C with 70% humidity while having access to sugar water and sterilized bee pollen ad libitum until sampling.

Except if otherwise mentioned, bacterial solutions were prepared on the inoculation day from fresh cultures grown in petri dishes for 16 to 20 hours. These cells were collected from petri dishes containing cells revived from −80°C stocks that were grown for 3-5 days in anaerobic conditions.

### Correlation between *F. perrara* and *Gilliamella* in bees colonized with the SynCom

Up to ten microbiota deprived honey bees per treatment were colonized with one of six different genotypes of *F. perrara* - wild type corresponding to strain PEB0191 - and a synthetic community (SynCom) composed of 13 members that represent the most abundant bacterial genera found in the honeybee gut (for more details please check (30)) that included 1 isolate of *S. alvi* and 3 isolates of *Gilliamella* – ESL169 (*Gilliamella* 53_1 also *Gilliamella apis*), ESL177 (*Gilliamella* 53_5) and ESL309 (*Gilliamella* 52_2, also *Gilliamella* sp.). In such experiment, *F. perrara* concentration, based on optical density, was 13 times that of each individual member of the SynCom. Bees were inoculated by handfeeding 5µl of a microbial solution containing simultaneously *F. perrara* and the SynCom members. Ileum samples were dissected at day 10 post bacterial inoculation. To identify SynCom members negatively correlated with *F. perrara,* absolute bacterial abundances were calculated using a) a targeted approach relying on qPCR for specific bacterial genera and b) an unbiased approach combining 16S amplicon sequencing with qPCR quantification using universal 16S primers. For the second approach, total bacterial abundance was calculated by multiplying the relative abundances for the 13 ASVs found within a given sample by its total number of 16S rRNA copies, obtained by qPCR using universal 16S rRNA primers. Correlations for bacterial absolute abundances were obtained using the function “*cor”* implemented in RStudio. Correlation coefficient values for all combinations can be found in the **Supplementary Dataset 1.**

### DNA extraction for qPCR bacterial quantification

Dissected ileums were homogenized with 165 µl of a solution containing GI lysis buffer, QIAGEN, and lysozyme (10:1 concentration) zirconia beads (0.1 mm dia. Zirconia/Silica beads; Carl Roth) and glass beads in a Fast-Prep24 5G homogenizer (MP Biomedicals) at 6 m/s for 45s. After homogenization, samples were incubated at 37°C for 30 min. Then, 30 µl of Proteinase K were added and samples were incubated at 56°C for 1 hr. The purification of nucleic acids was performed using CleanNGS magnetic beads (CNGS-0005) and the Opentron OT-2 pipetting robot. Purified DNA extracts were stored at –20°C until further use.

### Bacterial quantification by qPCR

Bacterial absolute abundances were determined using quantitative PCR (qPCR) assays targeting the 16S rRNA gene of *Frischella*, *Gilliamella* and *Snodgrassella* or relying on universal primers for the 16S rRNA gene. Normalization was based on the number of host actin gene copies as described in (45). Primer sequences are given in **Table S2**. qPCR was conducted on a QuantStudio5 instrument (Applied Biosystems) with the following run method: a holding stage consisting of 2 min at 50°C followed by 2 min at 95°C, 40 cycles of 15 s at 95°C, and 1 min at 60°C. A melting curve was generated after each run (15 s at 95°C, 20 s at 60°C and increments of 0.3°C until reaching 95°C for 15 s) and used to assess specificity of PCR products. qPCR reactions were performed in 10 µl reactions in triplicates in 384-well plates, and each reaction consisted of 1 µl of DNA, 0.2 µM of forward and reverse primer and 1x SYBR green ‘Select’ master mix (Applied Biosystems). Each DNA sample was screened with at least one bacterial primers targeting the 16S rRNA gene of *Frischella*, *Gilliamella*, *Snodgrassella* or universal 16S rRNA primer and the actin gene of *A. mellifera*. For each target, standard curves were generated for absolute quantification using serial dilutions (from 10^7^ to 10 copies) of the target amplicon cloned into the plasmid vector. Absolute abundance of bacteria was calculated using the standard curve and was normalized by the median actin copy number per condition (to account for differences in gut size) and by the amount of 16S rRNA copies per genome. For the calculation, we used the following formula:

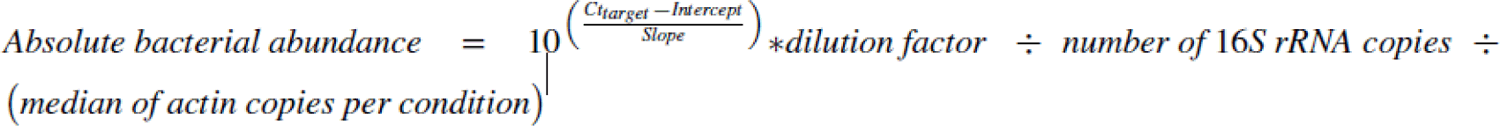

### 16S rRNA gene amplicon-sequencing

The V4 region of the 16S rRNA gene was amplified following the Illumina 16S metagenomic sequencing preparation guide and the protocols reported in ref (45, 46). We used primers 515F-Nex and 806R-Nex **(Table S2)**, to amplify the V4 region of the 16S rRNA gene. The PCR cocktail consisted of 12.5 μl of Invitrogen Platinum SuperFi DNA Polymerase Master Mix, 5 μl of MilliQ water, 2.5 μl of each primer (5 μM) and 2.5 μl of template DNA. PCR conditions were: 98 °C for 30 s followed by 25 cycles of 98 °C for 10 s, 55 °C for 20 s and 72 °C for 20 s and by a final extension step at 72 °C for 5 min. We verified that amplifications were successful by 2% agarose gel electrophoresis. We next purified the PCR products using Clean NGS purification beads (CleanNA) in a 1:0.8 ratio of PCR product to beads and eluted the purified PCR products in 27.5 μl of 10 mM Tris, pH 8.5. A second PCR step was performed to add a unique dual-index combination to each sample using the Nextera XT index kit (Illumina). We performed second-step PCR amplifications in a total volume of 25 μl, using 2.5 μl of the PCR products, 12.5 μl of Invitrogen Platinum SuperFi DNA Polymerase Master Mix, 5 μl of MilliQ water and 2.5 μl each of Nextera XT index primers 1 and 2. PCR conditions consisted of an initial denaturation step at 95 °C for 3 min followed by eight cycles of 30 s at 95 °C, 30 s at 55 °C and 30 s at 72 °C and a final extension step at 72 °C for 5 min. PCR products were purified using Clean NGS purification beads in a 1:1.12 ratio of PCR product to beads and eluted in 27.5 μl of 10 mM Tris, pH 8.5. We quantified amplicon concentrations by PicoGreen and pooled the libraries in equimolar concentrations (with the exception of negative PCR controls and blank DNA extractions, which we pooled in equal volumes instead). We then verified that the final pool was of the right size using a Fragment Analyzer (Advanced Analytical). Sequencing was performed at the Genomic Technology Facility of the University of Lausanne on an Illumina MiSeq sequencer for 500 cycles, producing 2 × 250 base pair (bp) reads.

### Analyses of 16S rRNA gene amplicon-sequencing data from the ileum samples of gnotobiotic bees

We sequenced 16S rRNA gene amplicons from ileum samples, negative PCR controls and blank DNA extractions. We also included a mock community sample consisting of equal numbers of nine plasmids (pGEM-T Easy vector; Promega) containing eight 16S rRNA gene sequences from honeybee gut symbionts and one from *Escherichia coli*, which we used as an internal standard to verify consistency between MiSeq runs. Raw sequencing data were quality-controlled with FastQC (47) and primer sequences were removed with Cutadapt (48). We then continued the analysis using the Divisive Amplicon Denoising Algorithm 2 (DADA2) package v.1.20.0 in R (49). All functions were run using the recommended parameters except that at the filtering step we truncated the F and R reads after 232 and 231 bp, respectively. We then set randomize=TRUE and nbases=3e8 at the learnErrors step. We used the SILVA database (v.138) to classify the identified ASVs. To complement the taxonomic classification based on the SILVA database, sequence variants were further assigned to major phylotypes of the bee gut microbiota as previously defined in (45). Unclassified ASVs were removed with the ‘phyloseq’ package v.1.36.0 (50), using the ‘subset taxa’ function. We then used both the ‘prevalence’ and ‘frequency’ methods (method = ‘either’) in the R package ‘decontam’ v.1.12.0 (51) to identify and remove contaminants introduced during laboratory procedures, using the negative PCR controls and the blank samples as reference.

### Correlation analysis of wild-derived *Apis mellifera* gut microbiomes

Wild-derived hindguts (pylorus, ileum and rectum) of *Apis mellifera* females were collected from two hives in Switzerland (49 individuals) and two hives in Japan (24 individuals). Samples were processed using detailed protocols found in (31). These, include the sample processing, DNA extraction, and bioinformatic pipeline performed to obtain the relative abundance of a total of 72 different bacterial species that comprise the gut microbiome of these 73 individual honey bees. Posteriorly, for all samples, the total 16S rRNA gene copy number was assessed using qPCR with a set of universal 16 primers **(Supplementary Table S2)**. In order to calculate bacterial absolute abundances per sample, the relative abundance of each bacterium was multiplied by the corresponding total counts of 16S rRNA gene copies. SparCC (52), which infers correlation networks from compositional data was computed. Specifically, bootstrapped estimates of SparCC correlation coefficients, were obtained by employing the *sparccboot* function with R=1000 of the R package SpiecEasy (53). Correlation coefficient values for all combinations can be found in the **Supplementary Dataset 2.**

### Phylogeny of the *Gilliamella* genus

To determine the phylogenetic placement of *Gilliamella* 52_2 and its relationship to other *Gilliamella* strains correlated with *F. perrara* in conventional bees, we reconstructed a genome-scale phylogeny of *Gilliamella* containing isolates from honey bees sampled across Japan, the USA, Switzerland, Sweden, and Australia, and including metagenome-assembled genomes from Japan and Switzerland ((31); **Supplementary Dataset 3**). Genome quality was assessed using CheckM v1.2.2, and only assemblies with >75% completeness and <10% contamination were retained for downstream analyses. Taxonomic assignments were performed with GTDB-Tk v2.4.1 using database release r226. Gene annotation was conducted with DRAM v1.4.6. Orthologous groups were identified with OrthoFinder v3.1.0, and single-copy orthologs were individually aligned and concatenated into a supermatrix using AMAS v1.0. Poorly aligned regions were removed with trimAl v1.5.0. Maximum-likelihood phylogenetic trees were inferred with IQ-TREE v3.0.1 under the LG+I+G4 substitution model, with 1000 bootstrap replicates. *F. perrara* strain PEB0191 was used to root the tree.

### Co-colonization of bees with *F. perrara* and *Gilliamella* sp. ESL309 at different ratios

On the day before honey bee inoculation, bacteria were harvested from three days old petri dishes, transferred to new solid media plates using a sterile plastic loop and cultured for 16-20 hours in anaerobic conditions at 35°C. *F. perrara* cells were amplified in TYG plates and *Gilliamella sp.* ESL309 were transferred to BHI medium.

Microbiota deprived bees were inoculated with three different ratios of *F. perrara* and *Gilliamella* sp. ESL309: 10:1, 1:1 and 1:10. For generating the 10:1 condition, 100µl of *F. perrara* at OD_600_=1 were mixed with 100µl of *Gilliamella sp.* ESL309 at OD_600_=0.1 and 800µl of a sugar water:PBS solution (SW:PBS). Following the same logic, 1:1 ratio was obtained by mixing 100µl of *F. perrara* at OD_600_=0.1, 100µl of *Gilliamella sp.* ESL309 at OD_600_=0.1 and 800µl of SW:PBS. The 1:10 ratio consisted of 100µl of *F. perrara* at OD_600_=0.1, 100µl of *Gilliamella sp.* ESL309 at OD_600_=1 and 800µl of SW:PBS. As controls for bacterial competition, honey bees were mono inoculated with *F. perrara* or *Gilliamella sp.* ESL309 at OD_600_=1 or OD_600_=0.1. By plating bacteria at defined OD_600_, we ensured that OD_600_ and CFU are equivalent in both species (see **Supplementary Figure S13**). At day 10 post-inoculation, up to 10 bee guts were dissected, the ileum portion was isolated, placed in a microcentrifuge tube, snap-frozen using liquid nitrogen and kept at −80°C until DNA extraction was performed. For each sample, bacterial loads were quantified by qPCR using *Frischella* and *Gilliamella* genus specific primers (**Supplementary Table S1**) normalized by the values for the honey bee actin gene, as previously described.

In this experiment, two different genotypes of *F. perrara* were tested, either wild-type or a mutant version lacking the hcp protein of the T6SS-2 (generated for (30)). A total of two different independent experimental replicates were performed. Per experiment and per treatment, a single plastic cup was used that contained up to 20 honey bees.

### Lifespan experiments

On the day before honey bee inoculation, bacteria were harvested from three days old petri dishes, transferred to new solid media plates using a sterile plastic loop and cultured for 16-20 hours in anaerobic conditions at 35°C. *F. perrara* cells were amplified in TYG plates and *Gilliamella sp.* ESL309 were transferred to BHI medium.

For the first set of experiments, 0-3 days old microbiota deprived adult bees were inoculated with *F. perrara*, *Gilliamella* sp. ESL0309, both bacteria simultaneously to understand the impact of competition on host fitness, or no bacteria. Each bacterium was inoculated at a final OD_600_=0.1 and each bee received 5µl of a bacterial solution diluted in PBS and sugar water at equal proportions. After inoculation and for each different condition tested, bees were split into three different cup cages (7-10 individuals) and provided pollen and sugar water *ad libitum*.

For the second set of experiments, 0-3 days old microbiota deprived adult bees were inoculated with *F. perrara*, *Gilliamella* sp. ESL0309 or no bacteria. Each bacterium was inoculated at a final OD_600_=0.1 and each bee received 5µl of a bacterial solution diluted in PBS and sugar water at equal proportions. At day 5 after the inoculation of the native microbiome, half of the honey bees were inoculated with 5µl of a bacterial solution of PBS and sugar water containing *Serratia ureilytica* at a final OD_600_=1 while the other half received the solution without bacterium. Survival was checked daily and dead bees were removed from the cage to minimize contaminations to the other honey bees. Two different independent experimental replicates were performed. Data was analyzed on R using proportional hazards models, specifically, the *coxme* function of package coxme was employed with experimental replicate as a random factor (54). For the infection experiment, the model included the interaction between microbiome composition (*F. perrara*, *Gilliamella* sp. ESL309 or no bacteria) and infection status. For both models, comparisons across treatments were performed using the emmeans package on the coxme model.

### Bacterial quantification of gut bacteria location within the ileum

Microbiota deprived bees were individually inoculated with 5µl of a bacterial solution containing *F. perrara* or *Gilliamella sp.* ESL309 at a final OD_600_=0.1. At day 7 post-inoculation the ileum portion of the gut was isolated and separated into anterior and posterior sections. To do so, a cut was performed using a dissection blade upstream of the location where the ileum naturally twists. Uncut ileums were kept as controls. Samples were placed in microcentrifuge tubes containing 1mL of PBS and glass beads and homogenized using Fast-Prep24 5G homogenizer (MP Biomedicals) at 6 m/s for 45s. Bacterial solutions were diluted in a 10-fold series in 96 well plates filled with PBS and bacterial loads were quantified by plating 5µl of each dilution into BHI plates. Plates were incubated for 3 to 5 days in anaerobic conditions at 35°C before CFU quantification. Two independent experimental replicates were performed and, per experiment, the loads of up to 10 guts per treatment were quantified.

### Generation of fluorescently tagged bacterial strains

To enable visualization of bacteria within the honey bee gut, fluorescently tagged strains of *F. perrara* and *Gilliamella* sp. ESL309 were generated. To accomplish that, wildtype strains of *F. perrara* or *Gilliamella* sp. were conjugated with *Escherichia coli* JKE201 auxotrophic strain for DAP, carrying a plasmid encoding either green fluorescent protein (pAC08 for *F. perrara*, Addgene #197402) or E2-Crimson (pAC09 for *Gilliamella* sp., Addgene #197403) under the control of a CP25 promoter (55). Both plasmids contain a RSF1010 origin of replication and an ampicillin resistance gene. Briefly, *F. perrara* or *Gilliamella* sp. were co-cultured with *E. coli* JKE201 carrying the plasmid of interest overnight on TSA medium supplemented with DAP to ensure conjugation. After conjugation, cells were washed and plated on TSA medium without DAP but supplemented with ampicillin (30 µg/mL) to select for successful transconjugants. Colony PCR was performed on the resulting colonies to amplify the plasmid gene encoding the fluorescent protein, confirming successful conjugation, (forward primer: AC06, reverse primer: AC08, **Table S2**). Moreover, PCR products were sequenced to confirm the presence of the fluorescent marker, and positive strains were stored in glycerol at −80 °C.

### Fluorescence microscopy imaging

Microbiota deprived bees were individually inoculated with 5 µl of a bacterial solution containing *F. perrara* carrying a plasmid expressing GFP or *Gilliamella sp.* ESL309 carrying a plasmid expressing Crimson, at a final OD_600_=1. At day 7 post-inoculation the anterior portion of the ileum was isolated for image acquisition. Contrarily to other experiments, in such context honey bees were not fed pollen, as pollen grains are auto fluorescent and interfere with image acquisition. Depending on the experiment, samples were either fixed overnight under gentle rotation at 4 °C in a 4% paraformaldehyde (PFA) solution, or imaged on the same day. Fixed samples were washed in the morning after fixation in PBS three times for 10 minutes each, with shaking at 4 °C. Before image acquisition, all samples were stained with 4′,6-diamidino-2-phenylindole (DAPI). Staining was performed by adding 1 µL of DAPI stock solution (5 µg/mL) to 1 mL of 0.5% Triton X-100 in PBS. Ileums were stained for 45 minutes to 1 hour under gentle rotation at 4 °C. After staining, samples were washed in PBS three times for 10 minutes each, under the same conditions. All samples were mounted on microscopy slides using nail polish cages containing 50 µL of PBS and covered with a microscopy slide.

For image acquisition, we utilized two different confocal microscopes, a ZEISS LSM 910 *c*onfocal microscope and a Leica Stellaris 8 microscope. In order to process a large scale of samples within the same day, a Zeiss LSM910 microscope was employed. GFP, Crimson and DAPI were excited at 488 nm (0.75% intensity), 561 nm and 405 nm (1% intensity) using a 43 μm pinhole, respectively. A 5× objective was used, 15 Z stacks were taken for each image as well as 4 or 9 tiles depending on the size of the picture. If a signal suggesting bacterial colonization was detected, images were acquired with a 40x objective using the above-mentioned parameters. Samples imaged included ileums fixed overnight and ileums imaged immediately after dissection and staining with DAPI. Three guts colonized with *F. perrara-*GFP or *Gilliamella sp.* ESL309-Crimson were imaged and microbiota-deprived guts were used as negative controls. For three-dimensional imaging, guts were dissected to isolate the anterior ileum and were imaged on the same day. Confocal imaging was carried out using a Leica Stellaris 8 laser-scanning microscope equipped with a tuneable white-light laser (WLL) and a 405 nm diode laser. DAPI, GFP, and Crimson were excited at 405 nm, 488 nm, and 611 nm, respectively. Fluorescence emission was collected using detection windows of 430–484 nm for DAPI, 500–550 nm for GFP, and 616–834 nm for Crimson. Overview scans were obtained using a 10x objective, and z-stacks were acquired using 63x objective. In both cases, stacks were acquired as tiles and stitched using the LASX software.

## Statistical analyses and figure generation

Statistical analysis for the colonization data was performed using RStudio. For comparison between two conditions, Wilcoxon sum-rank test was employed as data was non-parametric. These comparisons include the effect of competition on colonization (absence versus present of the competitor) and the comparison across different portions of the ileum (anterior versus posterior versus full ileum).

Plots were generated on R using the *ggplot2* package (56). Microscopy images were analysed using ImageJ and the ZEN Microscopy Software. Phylogeny was generated using iTOL (57). The final figures were assembled using Affinity Designer 2.

## Data availability

The raw data and scripts used to generate the analyses and figures here presented can be found with the link <u>SantosMatos_ISME2026</u>. Data for the 16S rRNA sequencing is found at BioProject PRJNA1421104.

## Acknowledgments

We thank the Lausanne Genomics Technology Facility (GTF) team at the University of Lausanne for performing the 16S Amplicon Sequencing for the experiments. This work was supported by the NCCR Microbiomes, a National Centre of Competence in Research funded by the Swiss National Science Foundation (SNSF, grant number 180575 and 225148, to P.E. and Y.S.), the SNSF Consolidator grant GLOBEE (grant number 213860, to P.E.), and a FORESTO Program (grant number JPMJFR201C, to R.M.), funded by Japan Science and Technology Agency.

## Supplementary Figures

**Supplementary Figure S1.**
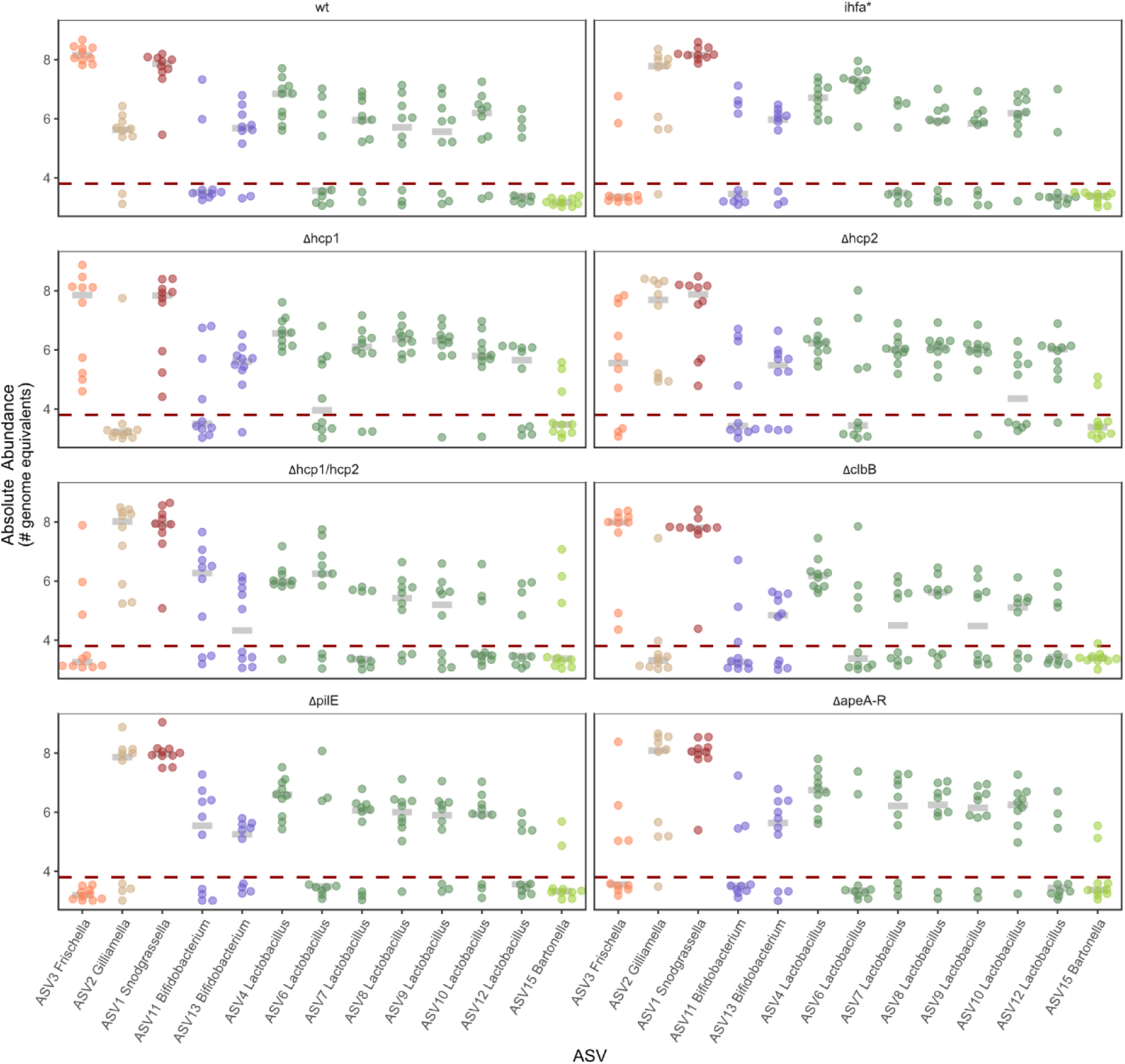
ASV abundance for the SynCom Experiment. Absolute Abundances (Y-axis) of the 13 ASV (X-axis) found by amplicon sequencing of 16S rRNA gene for the SynCom samples. Each colour depicts a different genus and each point represents a different bee sample. The grey lines show the median value for each ASV, while the dark red dotted line represents the limit of detection, based on the number of reads and the total bacterial load in each sample. Each panel represents a different *F. perrara* genotype tested: wt - wild type, ihfa* - spontaneous mutant for the subunit A of the transcription factor IHF, Δhcp1 – deletion mutant for the hcp gene of the T6SS-1, Δhcp2 – deletion mutant for the hcp gene of the T6SS-2, Δhcp1/ Δhcp2 – deletion mutant for the hcp genes of the T6SS-1 and T6SS-2, ΔclbB - deletion mutant for the colibactin synthesis pathway, ΔpilE – deletion mutant lacking pili, ΔapeA-R – deletion mutant lacking the APE biosynthesis cluster. For more information on the mutants, check (30).

**Supplementary Figure S2.**
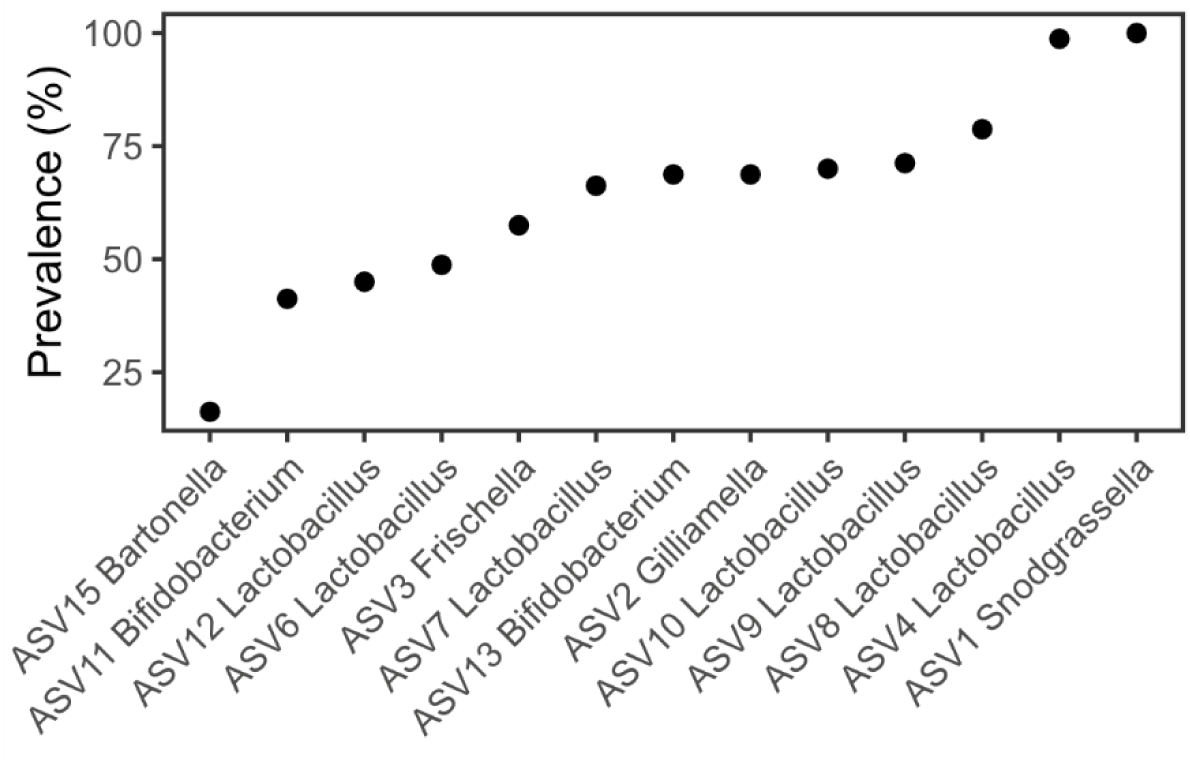
ASV prevalence across all samples of the SynCom experiment. Prevalence (Y-axis) of the 13 ASVs detected across the 80 bees samples of the SynCom experiment, for which 16S rRNA gene amplicon sequencing data was generated.

**Supplementary Figure S3.**
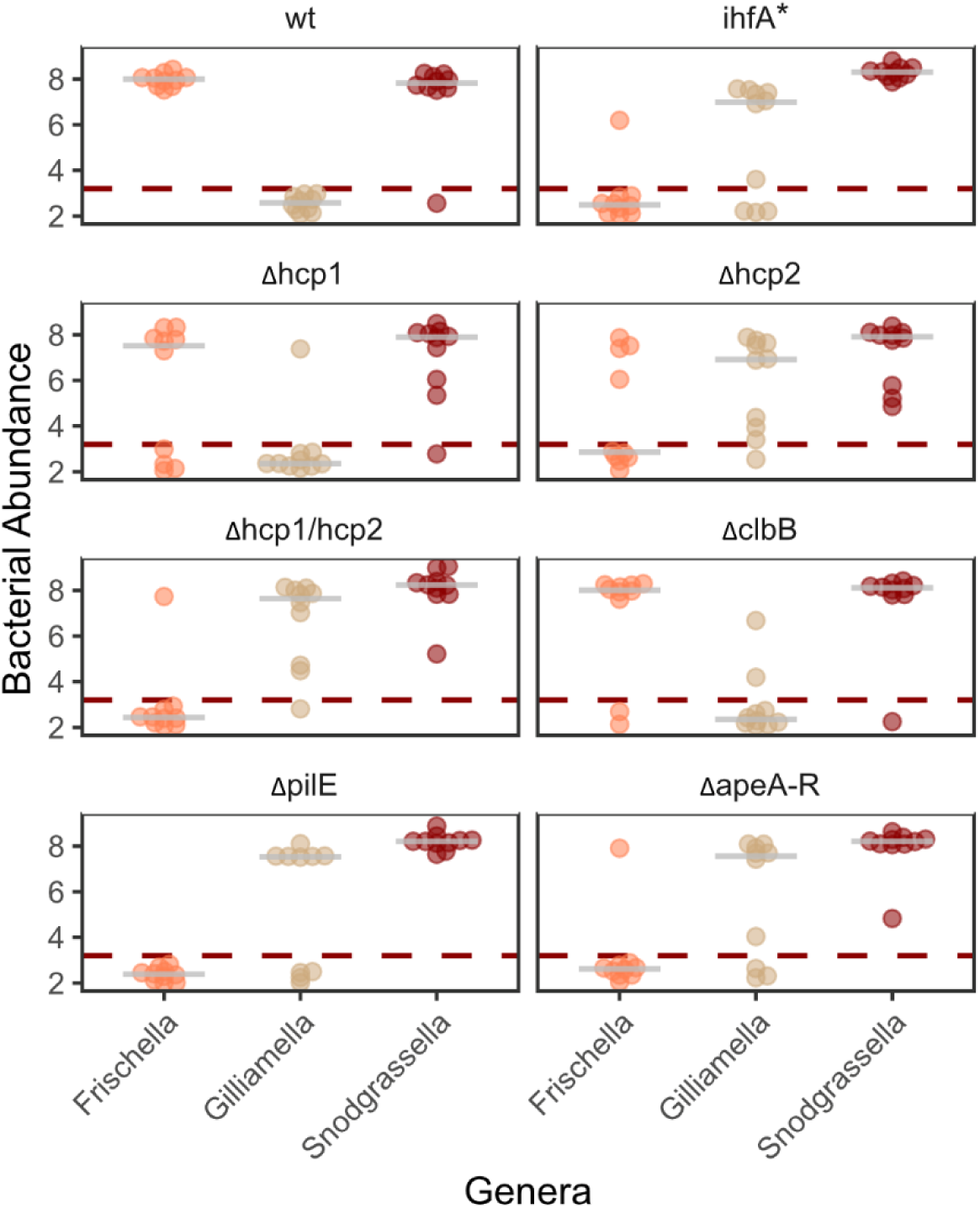
*Frischella* and *Gilliamella* do rarely co-occur in bee samples from the SynCom experiment. Absolute abundances (Y-axis) of *Frischella*, *Gilliamella* and *Snodgrassella* (X-axis) based on qPCR with genus specific primers, obtained for the bee samples of the SynCom experiment. Each color depicts a different ASV (orange: *Frischella*, beige; *Gilliamella* and brown: *Snodgrassella*) and each point represents a different bee sample. The grey lines show the median value for each ASV, while the dark red dotted line represents the limit of detection. Each panel shows a different *F. perrara* genotypes tested. For more information on the mutants, check (30).

**Supplementary Figure S4.**
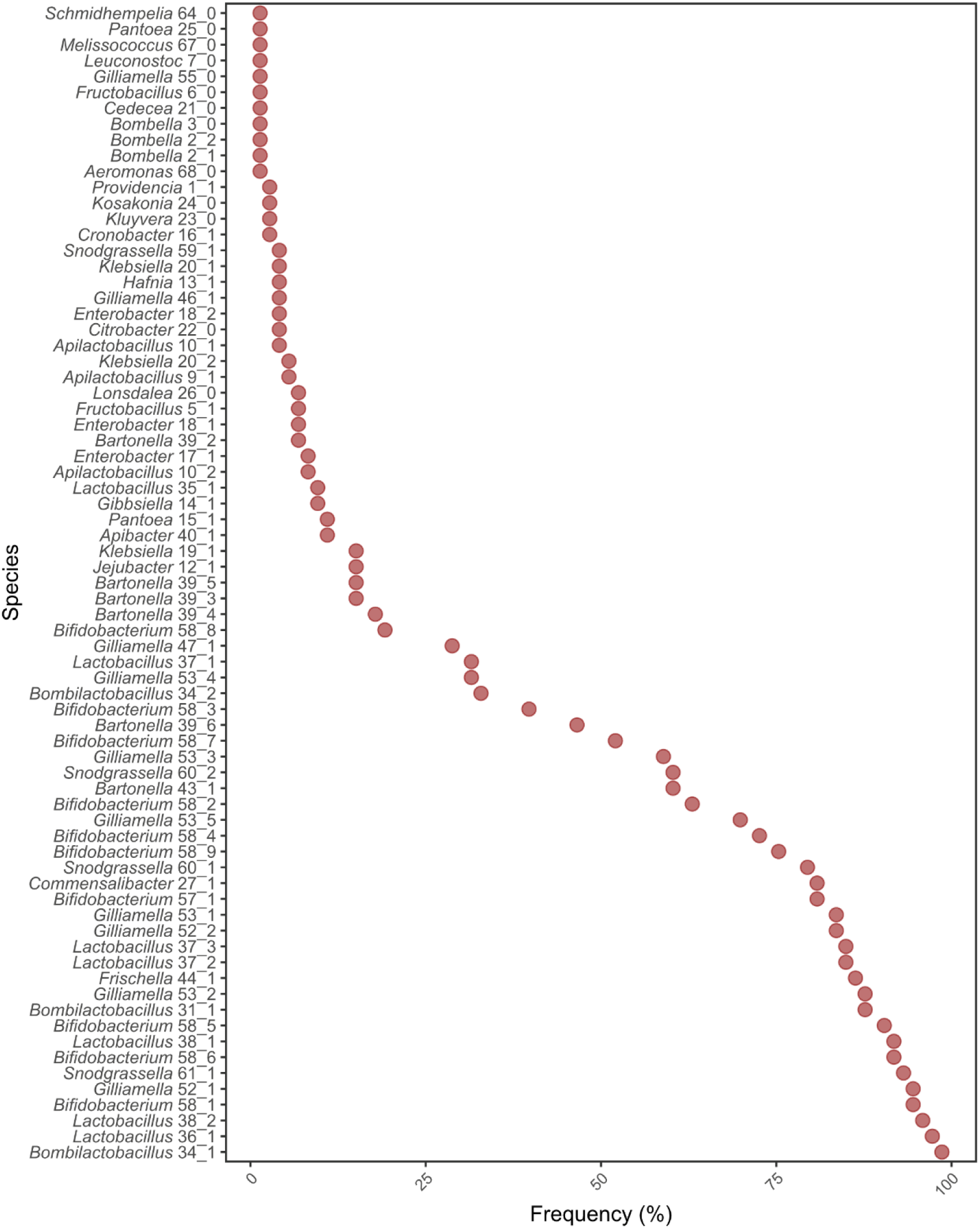
Species prevalence within the gut of conventional *A. mellifera*. Prevalence (X-axis) of the 72 bacterial species identified by metagenomic analysis of the gut microbiota of 73 honey bee samples from Switzerland and Japan. Species were identified by clustering metagenome-assembled genomes into species-level genome bins (see methods).

**Supplementary Figure S5.**
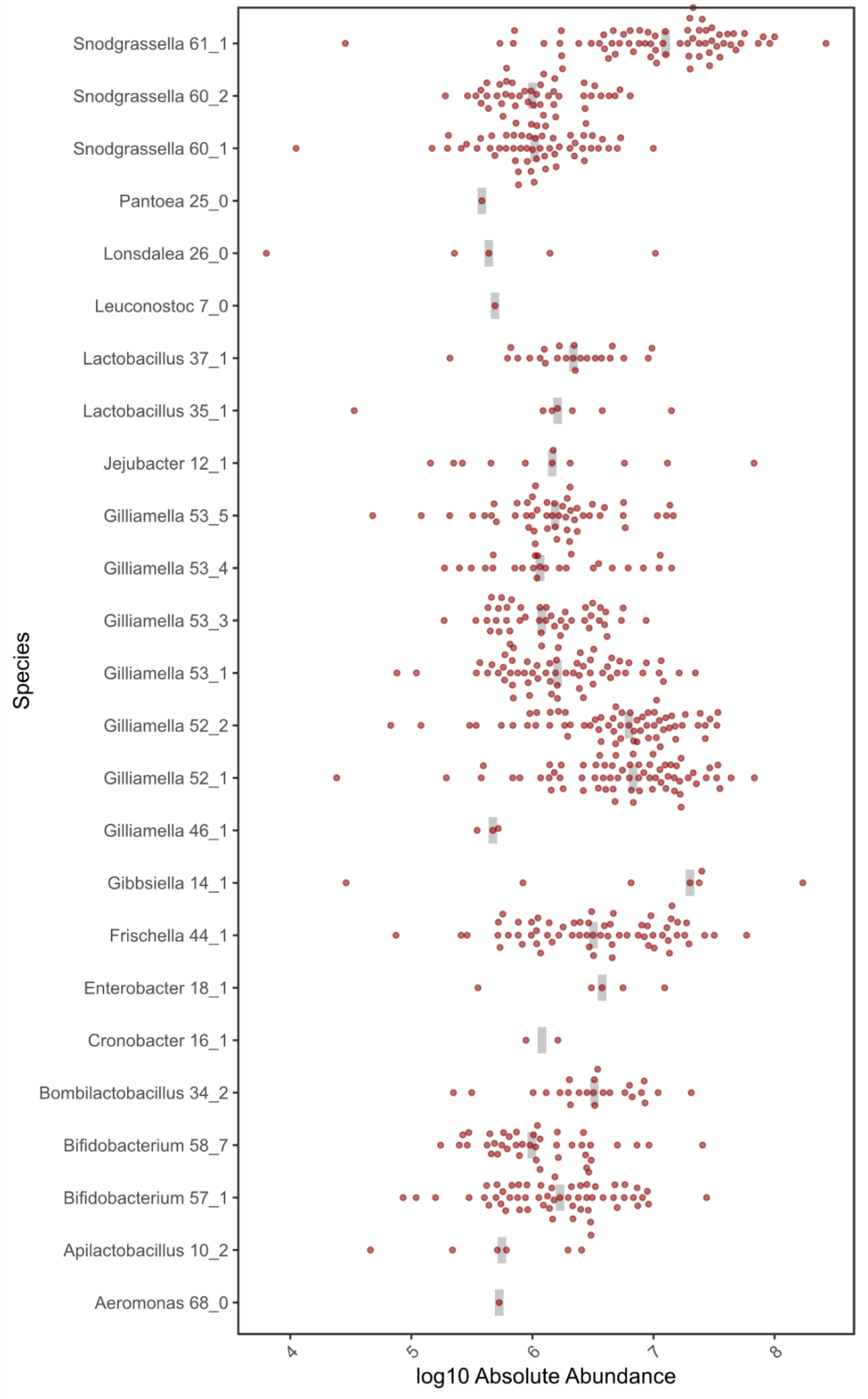
Absolute abundances of the 24 species significantly correlated with *F. perrara* within the gut of conventional honey bees. Absolute abundances (X-axis) for the 24 bacteria (Y-axis) shown in Figure 1B that were statistically significantly correlated with *F. perrara*. Each point depicts a different bee sample and only samples with values above the detection limit were plotted. Grey lines show the median value for each species.

**Supplementary Figure S6.**
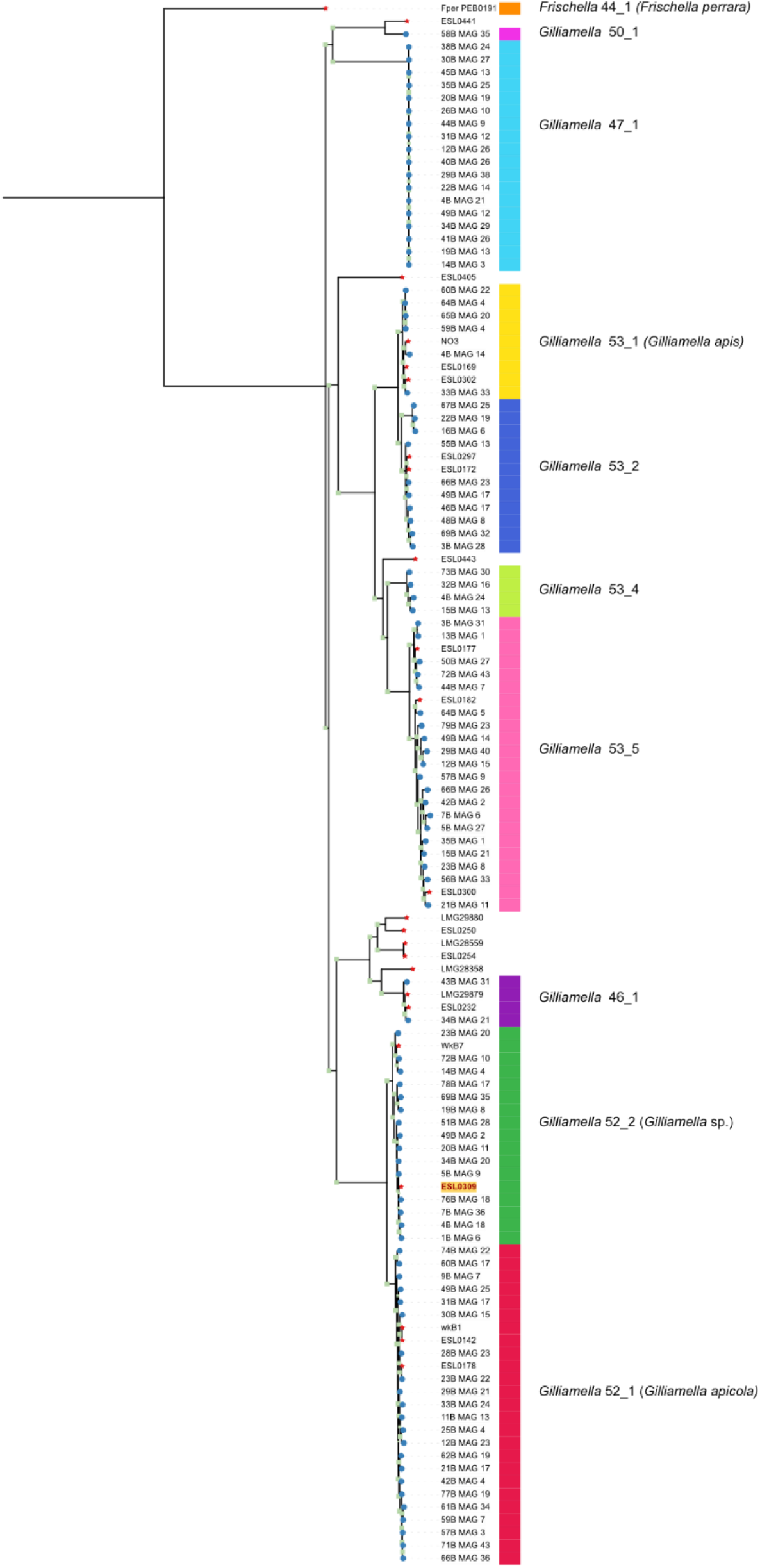
Phylogeny of the *Gilliamella* genus. Maximum-likelihood phylogeny of the *Gilliamella* genus based on the concatenated alignment of the protein sequences of xxx genes of 122 bacterial genomes (24 isolates and 98 metagenome-assembled genomes). These include 98 metagenome-assembled genomes, represented by the blue circles, and 24 genomes of bacterial isolates, represented with red stars. Bootstrap values above 90 (out of 100 replicates) are depicted with light green squares. The LG+I+G4 substitution model with 1000 bootstrap replicates was used to calculate the phylogeny. Species names shown by the different coloured rectangles correspond to those identified in the metagenomics approach. ESL 309, belonging to the species-level genome bin *Gilliamella* 52_2. is highlighted in dark red. *F. perrara* was used as an outgroup to root the tree. Note that some isolates included in the tree were not detected in the analyzed metagenomes and hence did not receive a species-level genome bin name.

**Supplementary Figure S7.**
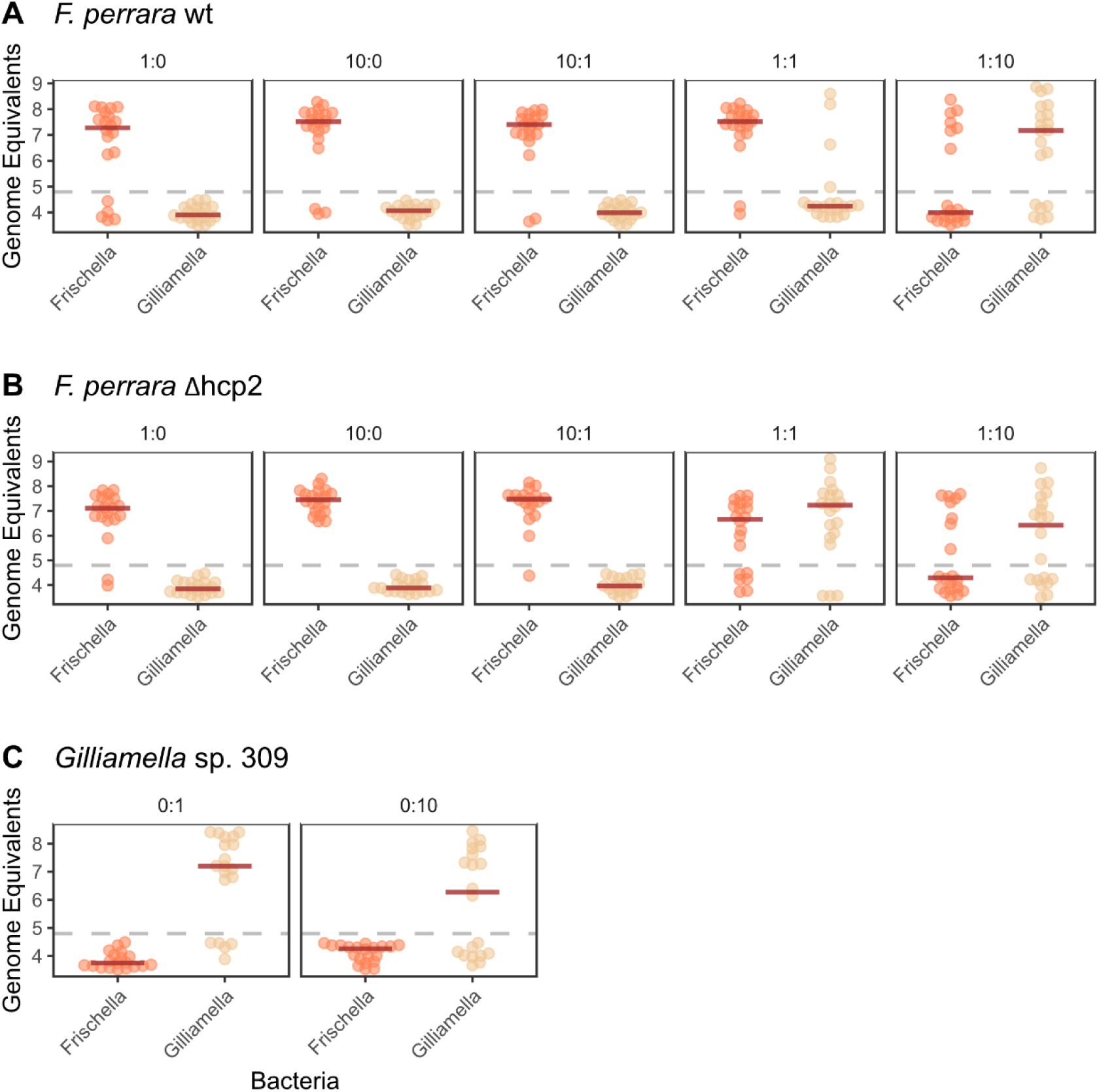
Colonization of *F. perrara* and *Gilliamella sp.* ESL309 across the different bacterial ratios tested. Bees were inoculated with *F. perrara* alone – either wt or Δhcp2 - (1:0, 10:0), *Gilliamella* sp. 309 alone (0:1, 0:10) or a combination of one *F. perrara* genotype and the *Gilliamella* strain at different ratios (10:1, 10:10 and 1:10). Bacterial loads, shown in the Y-axis, were quantified at day 10 post colonization using qPCR genus specific primers and normalized to the host actin levels for each sample. For each bacterium genus, depicted in the X-axis, each dot represents a different honey bee sample. Two experimental replicates were performed. The values depicted here were used to calculate the log2 fold differences between colonization alone and colonization in competition, used to generate Figure 2. For the ratio nomenclature, the number 10 corresponds to a bacterial solution at a final OD_600_ = 0.1, the number 1 to a final OD_600_ = 0.01 and 0 corresponds to the absence of bacterium. The first number refers to *F. perrara*, while the second one refers to *Gilliamella* sp. ESL309. **A** - Colonization levels for the ratios with *F. perrara* wt; **B** - Colonization levels for the ratios with *F. perrara* Δhcp2; C – Colonization levels of *Gilliamella* sp. ESL309 alone

**Supplementary Figure S8.**
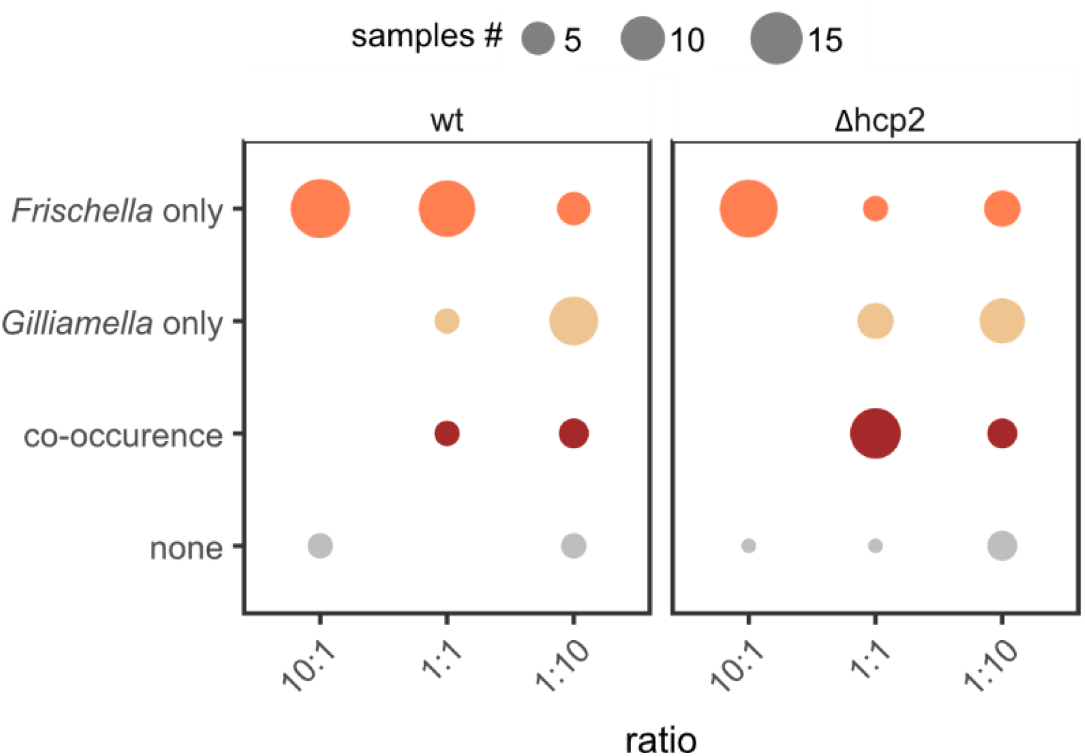
Inoculation size and the T6SS-2 of *F. perrara* underly inter-individual variation for colonization. Bacterial occurrence (Y-axis) was measured for each sample across the three ratios tested (X-axis), for the competition between *Gilliamella* sp. ESL309 against the wt or the Δhcp2 of *F. perrara*. If a given bacterium had levels above the limit of detection, it was considered as a colonizer of that gut sample. The size of the sphere shows the number of samples within each category. Data analysed here corresponds to the samples also depicted in Figure 2 of the main text.

**Supplementary Figure S9.**
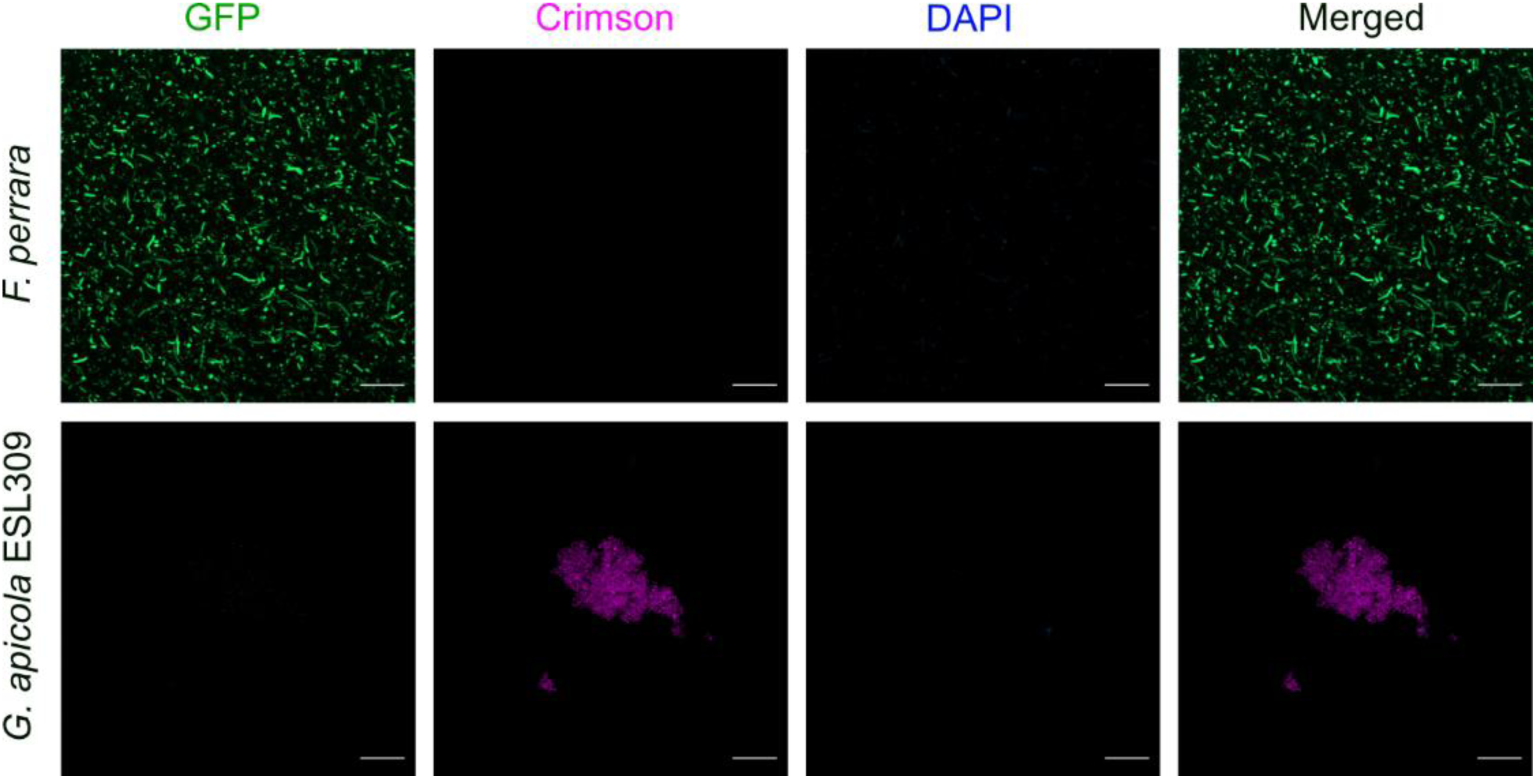
Visualization of *F. perrara*-GFP (green) and *Gilliamella* sp. ESL309 (magenta) *in vitro*. Scale bar corresponds to 20 µm. Images were obtained using a Leica Zeiss LSM900 confocal microscope with the 40x objective. Cells were imaged in the DAPI channel as a control, even though there was no DAPI labelling. Cells were grown in plates overnight prior to image acquisition.

**Supplementary Figure S10.**
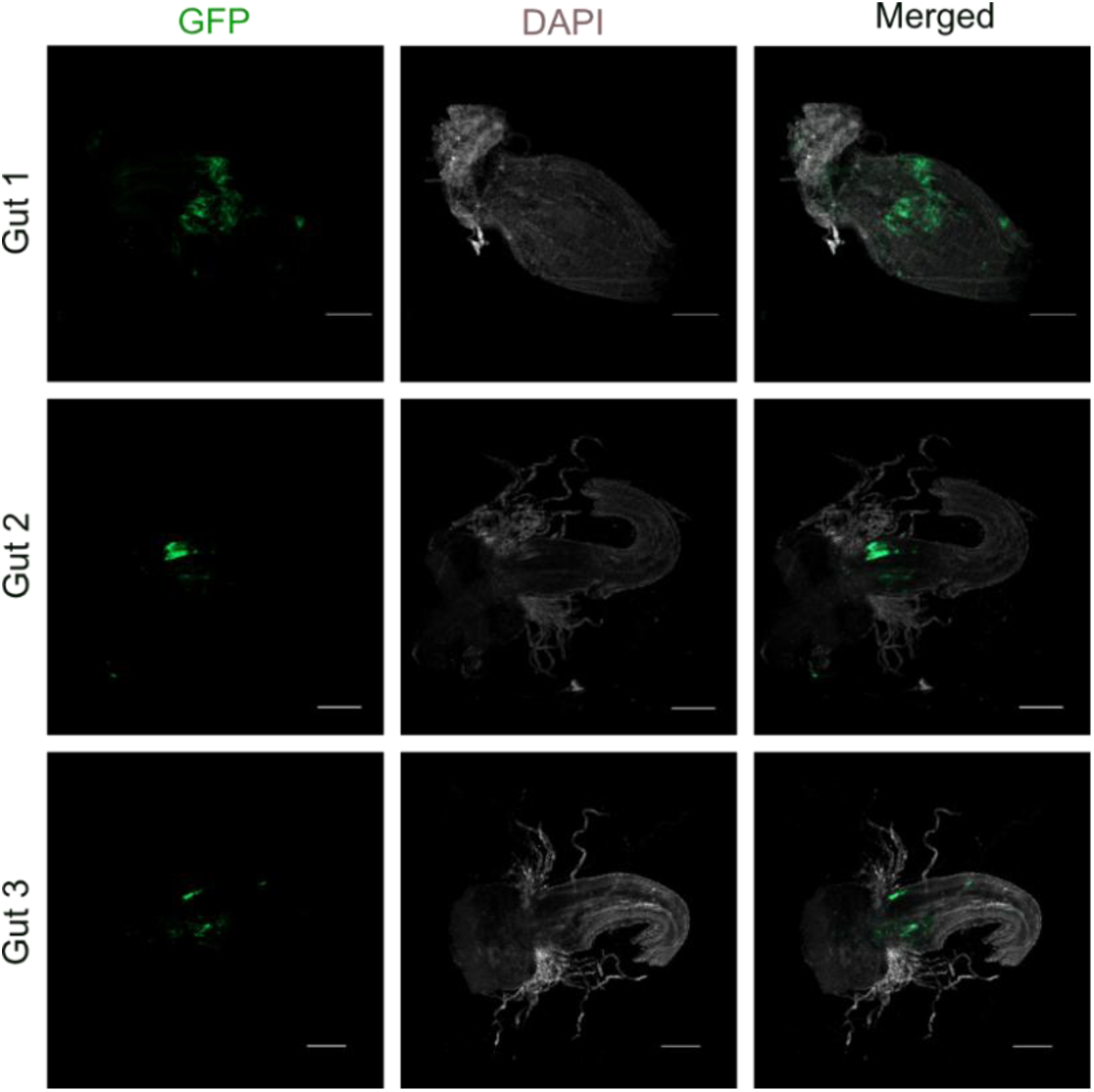
Visualization of the colonization of *F. perrara*-GFP (green) within the anterior hindgut of *A. mellifera*. Scale bar corresponds to 500 µm. Per gut, several images were obtained (tiles) using a Leica Zeiss LSM900 confocal microscope with the 5x objective and reconstructed relying on the ZEN3.1 software.

**Supplementary Figure S11.**
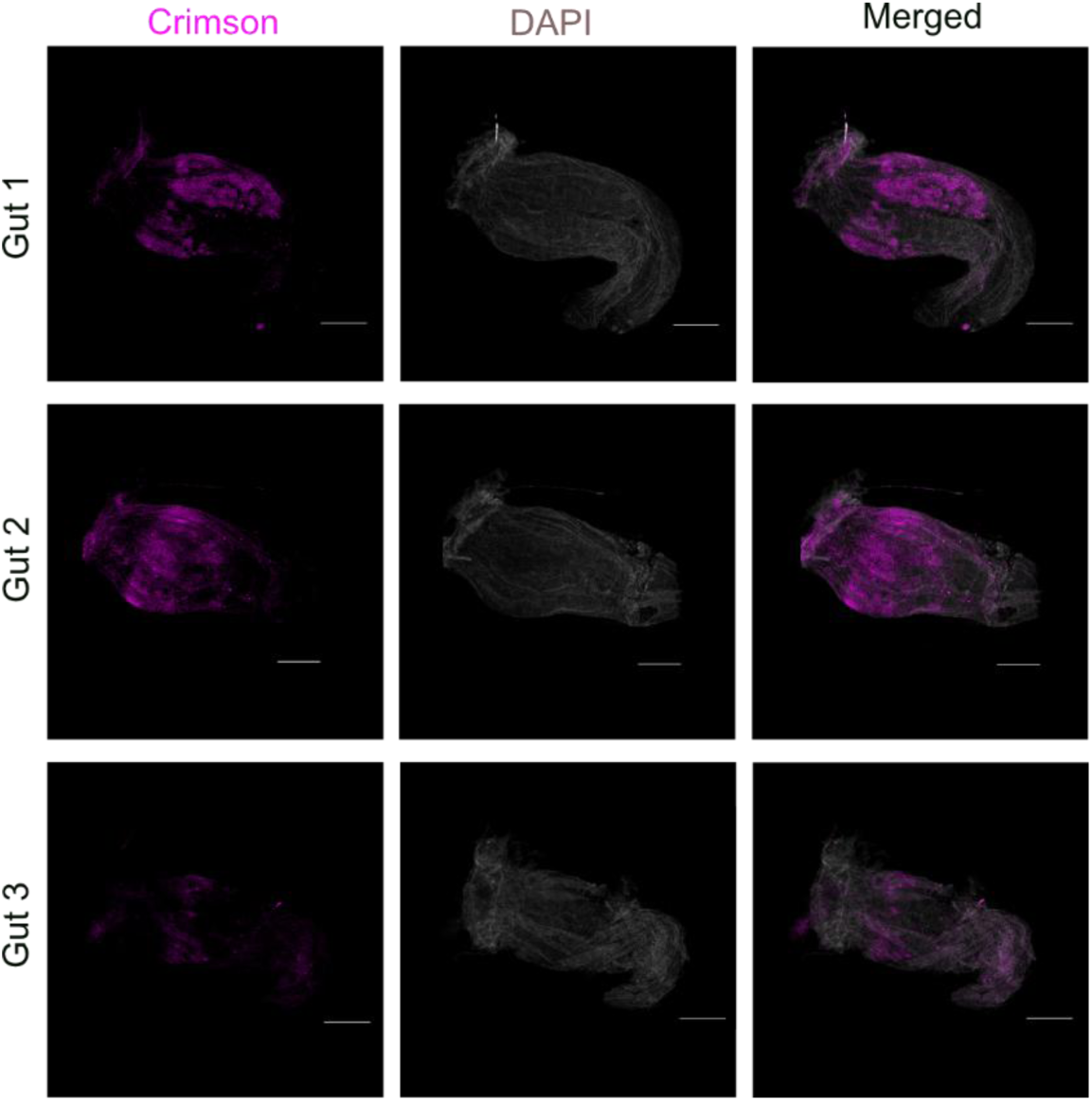
Visualization of the colonization of *Gilliamella* sp. ESL309 (magenta) within the anterior hindgut of *A. mellifera*. Scale bar corresponds to 500 µm. Per gut, several images were obtained (tiles) using a Leica Zeiss LSM900 confocal microscope with the 5x objective and reconstructed relying on the ZEN3.1 software.

**Supplementary Figure S12.**
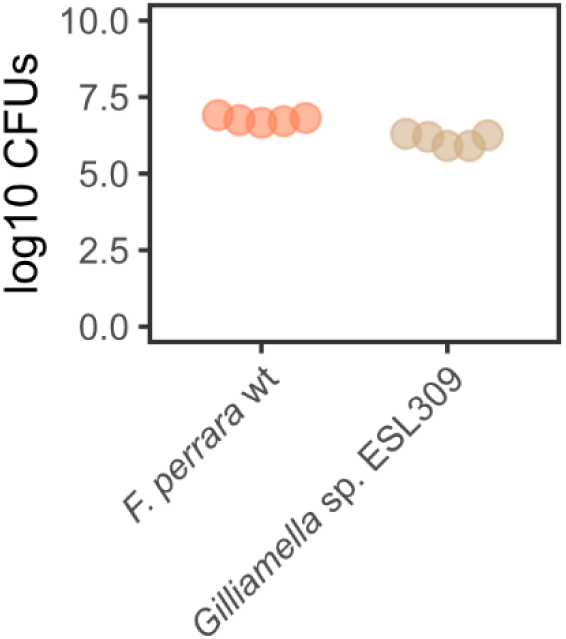
Correspondence between OD_600_ and CFU. Colony forming units (Y-axis) within 5µl of a *F. perrara* wt and *Gilliamella sp.* ESL309 solution at OD_600_=1. Bacterial solutions were plated in BHI petri dishes and grown for three days before quantification

## Supplementary Tables

**Supplementary Table S1.** Bacterial isolates used in this study.

| Species | Genotype | Strain ID | Reference | Experiment |
| --- | --- | --- | --- | --- |
| *Frischella perrara* | wild-type | ESL0868 | (25) | Ratio, Location, Intra-species competition, Lifespan and Survival after infection |
| *Frischella perrara* | Δhcp2 | ESL0855 | (30) | Ratio, Intra-species competition |
| *Frischella perrara* | wild-type, pAC08-GFP | ESL1281 | this study | Location |
| *Gilliamella* sp. | wild-type | ESL0309 | this study | Ratio, Location, Lifespan and Survival after infection |
| *Gilliamella* sp. | wild-type, pAC09-Crimson | ESL1283 | this study | Location |
| *Serratia ureilytica* | wild-type | ESL0497 | (58) | Survival after infection |

**Supplementary Table S2.**
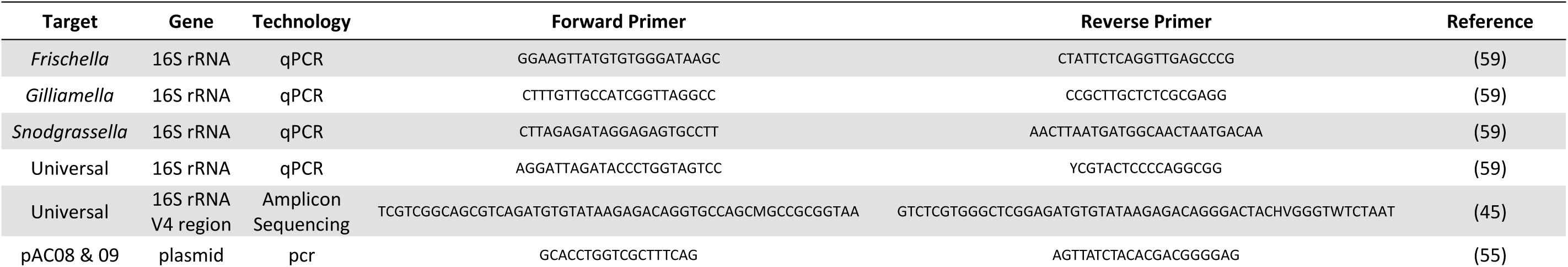
List of Primers used in this study.

